# The singularity of Cetacea behaviour parallels the complete inactivation of melatonin gene modules

**DOI:** 10.1101/490219

**Authors:** Mónica Lopes-Marques, Raquel Ruivo, Luís Q. Alves, Nelson Sousa, André M. Machado, L. Filipe C. Castro

## Abstract

Melatonin, the *hormone of darkness*, is a peculiar molecule found in most living organisms. Emerging as a potent broad-spectrum antioxidant, melatonin was repurposed into extra roles such as the modulation of circadian and seasonal rhythmicity: affecting numerous aspects of physiology and behaviour, including sleep entrainment and locomotor activity. Interestingly, the pineal gland—the melatonin synthesising organ in vertebrates—was suggested to be absent or rudimentary in some mammalian lineages, including Cetacea. In Cetacea, pineal regression is paralleled by their unique bio-rhythmicity, as illustrated by the unihemispheric sleeping behaviour and long-term vigilance. Here, we examined the genes responsible for melatonin synthesis (*Aanat* and *Asmt*) and signalling (*Mtnr1a* and *Mtnr1b*) in 12 toothed and baleen whale genomes. Based on an ample genomic comparison, we deduce that melatonin-related gene modules are eroded in Cetacea.

## Introduction

Circadian rhythmicity is critical for broad organism homeostasis. In mammals, a complex network of brain anatomical structures, such as the suprachiasmatic nuclei (SCN) and the pineal gland, as well as a set of specific molecular pathways, optimize physiological timekeeping^1,2^. First described in the 1950s, melatonin was since suggested to act as multi-purpose hormone, with important roles in SCN excitability and sleep modulation^3–7^. In vertebrates, endogenous melatonin levels exhibit daily rhythmicity and a secretory peak at night (Figure 1). Melatonin is predominantly synthesized in the pineal gland, with the canonical synthesis pathway initiating from a precursor molecule, the amino acid tryptophan^3^. The two final steps of the enzymatic cascade are sequentially controlled by aralkylamine N-acetyltransferase (AANAT) and hydroxyindole-O-methyltransferase (ASMT) enzymes, respectively: converting the intermediate serotonin into melatonin^8,9^. Subsequent hormone signalling in target cells occurs *via* two high affinity G-protein couple receptors, (MTNR1A) and (MTNR1B)^10^. Interestingly, vertebrate AANAT functional evolution, towards the conversion of serotonin into the substrate of ASMT N-acetyl serotonin, was suggested to parallel the emergence of the vertebrate pineal gland and melatonin rhythms^11^. Yet, some mammalian lineages, including Cetacea, seem to lack a structured pineal gland^12,13^. In fact, the existence of a functional pineal gland in these aquatic mammals is still contentious, even among individuals of the same species: with conflicting reports from various whale species (e.g. *Megaptera novaeangliae* (humpback whale)) or the commonly studied *Tursiops truncatus* (bottlenose dolphin)^14^. Paradoxically, a report using bottlenose dolphins lacking a visible pineal gland suggests the existence of measurable amounts of circulating melatonin but without a clear circadian rhythmicity^14^. To further address the parallelism between pineal regression and melatonin synthesis and signalling, we extracted genomic information and addressed the coding status of *Aanat, Asmt* and the melatonin receptors *Mtnr1a* and *Mtnr1b* in Cetacea and in *Hippopotamus amphibius* (common hippopotamus), the closest extant lineage of Cetacea.

**Figure 1:**
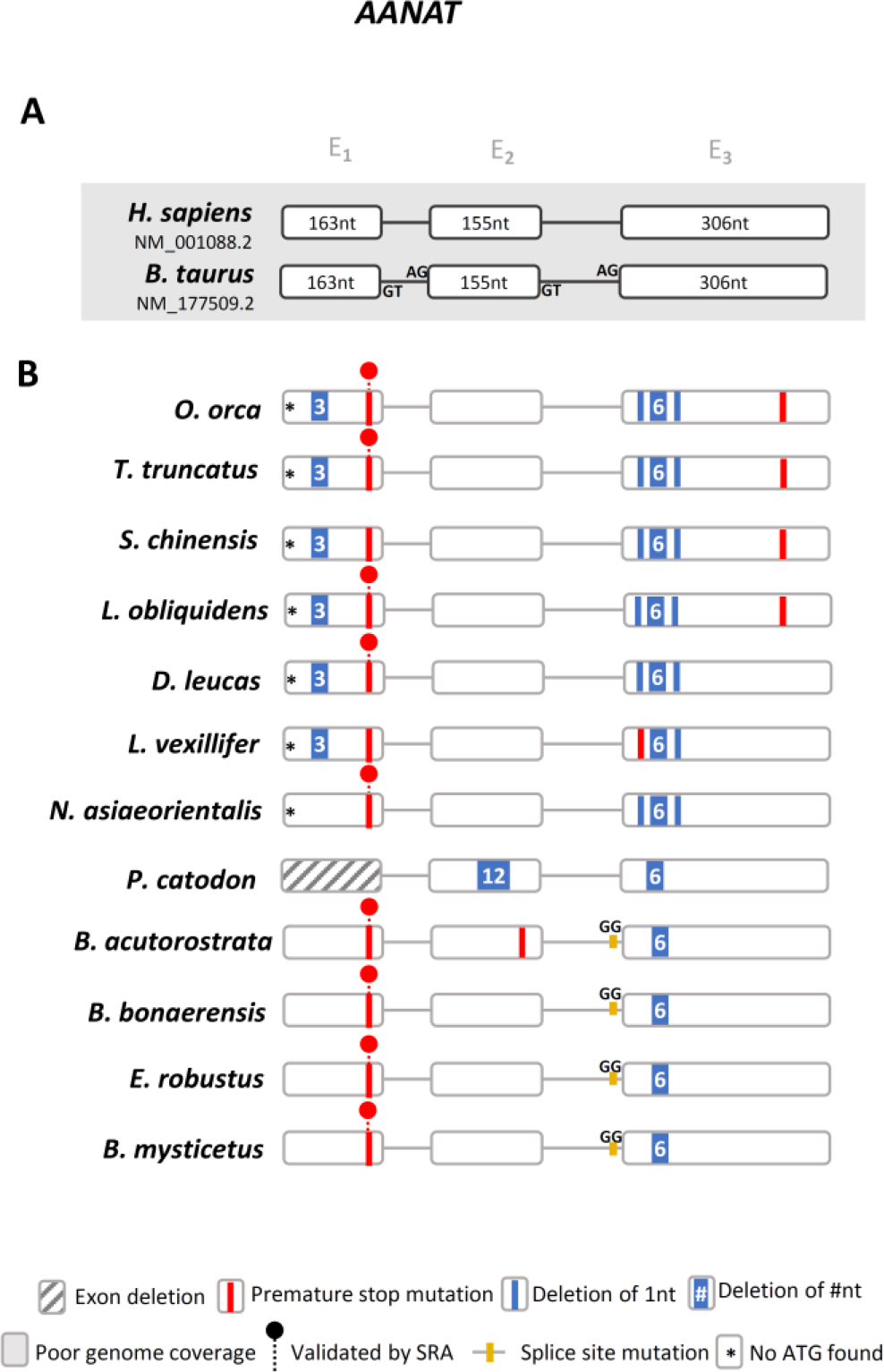
*Aanat* gene annotation. **A -** Schematic representation of the gene structure of human and *B. taurus Aanat* gene, each box represents an exon and lines represent intronic regions (not to scale). **B -** Schematic representation of the corresponding *Aanat* genes identified in Cetacea and location of the identified mutations. Non canonical splice sites are indicated above the corresponding annotation.

## Results

Using the available genomic data we examined the genomic regions of melatonin synthesis and signalling modules in 12 Cetacea species (*Orcinus orca* (orca whale), *Tursiops truncatus* (bottlenose dolphin), *Lagenorhynchus obliquidens* (Pacific white-sided dolphin), *Sousa chinensis* (Indo-Pacific humpbacked dolphin), *Delphinapterus leucas* (beluga whale), *Lipotes vexillifer* (Yangtze river dolphin), *Physeter catodon* (sperm whale), *Balaenoptera acutorostrata* (common minke whale), *Balaenoptera bonaerensis* (Antarctic minke whale), *Eschrichtius robustus* (grey whale), *Balaena mysticetus* (bowhead whale))^15–23^. We started by analysing the *Aanat* gene. The *Aanat “locus-of-origin”* was first identified using the human (NM_001088.2) and the *Bos taurus* (domestic cow; NM_177509.2) gene sequences (Supplementary Figure 1). In the case of *T. truncatus*, the *Aanat* gene was not annotated as a coding open reading frame (ORF) yet, the gene *locus* and corresponding neighbouring gene families were found to be conserved (Supplementary Figure 1). Next, we performed a comparative analysis of the exonic sequences, using the cow orthologue as reference, to identify potential mutations^24–27^. In summary, we were able to deduce the presence of multiple and shared disruptive mutations (Figure 1 and Supplementary Figure 2). These included a stop codon in exon 1 present in all analysed Cetacea with the exception of *P. catodon* (exon 1 was not found in the current assembly); an absent starting codon and frameshift deletions in exon 3 in Odontoceti (toothed whales); and a disruptive intron 2/exon 3 boundary mutation in Mysticeti (baleen whales). In *P. catodon* a unique deletion of 12 nucleotides in exon 2 was observed, which, combined with an additional 6 nucleotide deletion in exon 3, strongly suggests an inactivated ORF. In *O. orca* and *T. truncatus* a premature stop codon truncates exon 3. Other species-specific mutations were identified. To further confirm our findings, we selected the most critical and evolutionary conserved ORF-disruptive mutation for validation using Sequencing Read Archives (SRAs). As such, we confirmed the presence of the premature stop codon in exon 1 across 9 Cetacea species (Supplementary Figure 3). Regarding the Odontoceti *O. orca, T. truncatus, D. leucas, N. asiaeorientalis* and the Mysticeti *B. acutorostrata* and *E. robustus*, we were able to validate the conserved premature stop using two independent sequenced samples per species.

Next, we examined the *Asmt* gene (Supplementary Figure 4). In human, 3 distinct isoforms of *Asmt*, resulting from the alternative splicing of exon 6 and 7, were identified^28^. Yet, only isoform 1 (P46597) has been reported to display ASMT enzymatic activity (Supplementary Figure 5)^28^. Thus, both human and *B. taurus* genes were used as reference. Gene annotation revealed the presence of a conserved 1 nucleotide insertion in exon 1, validated by independent SRAs in *O. orca, T. truncatus, D. leucas, N. asiaeorientalis, B. acutorostrata* and *E. robustus* (Figure 2 and Supplementary Figures 6 and 7). In *L. obliquidens, S. chinensis, P. catodon* and *B. mysticetus* exon 1 could not be retrieved using the current assemblies. Complete erosion of exon 5 was also detected in *O. orca*, *L. obliquidens*, *S. chinensis*, *N. asiaeorientalis* and *D. leucas*, for which full genomic sequences ranging from exon 4 to exon 7 were available; for *P. catodon* and *E. robustus* exon 5 was identified with no deleterious mutations, while for the remaining species the presence of sequencing gaps within this region impaired further assessment. The deletion of exon 5 is expected to abolish enzyme activity given that this exon contains a highly conserved glycine (Gly187), participating in the active site of ASMT^28^ (Supplementary Figures 5 and 8). Also, aside from the conserved non canonical splice site observed in exon 8 of Odonoceti, other non-conserved ORF disrupting mutations were found (Figure 2): notably, a 1 nucleotide insertion in exon 4 of *P. catodon*, validated by independent SRAs (Supplementary Figure 9); and a 1 nucleotide deletion in exon 3 of *L. obliquidens*, also validated by SRA (Supplementary Figure 10). Finally, exon 9 was not found in the present genome assemblies of all analysed species (Figure 2).

**Figure 2:**
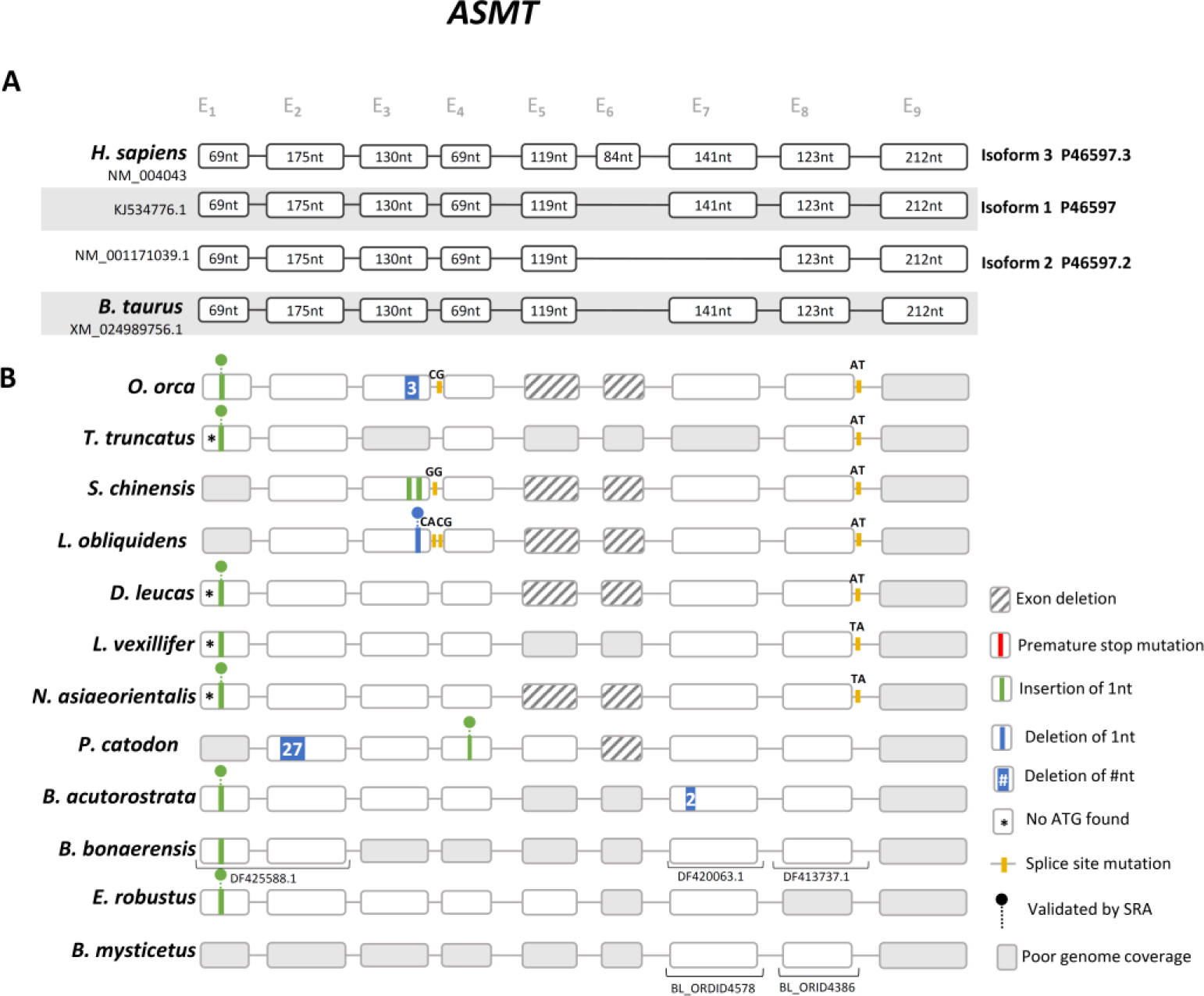
*Asmt* gene annotation. **A -** Schematic representation of the gene structure and distinct isoforms of human and *B. taurus Asmt* gene (not to scale), each box represents an exon and lines represent intronic regions. **B -** Schematic representation of the corresponding *Asmt* genes in Cetacea and location of the identified mutations. Non canonical splice sites are indicated above the corresponding annotation.

We next examined *Mtnr1a* and *Mtnr1b* genes, which are the receptors responsible for transducing melatonin signalling. Here, we failed to detect *Mtnr1a*-like sequences in Odontoceti genomes. Comparative synteny analysis revealed that *Fat1* and *F11*, which are adjacent to *Mtnr1a* in human and cow, are also side-by-side in Odontoceti (Figure 3). To further validate this observation, we carefully examined the genomic sequences flanked by *Fat1* and *F11 genes*. While in some species we observed the occurrence of sequencing gaps, impairing a robust conclusion regarding *Mtnr1a* absence, in *L. obliquidens, D. leucas* and *N. asiaeorientalis* the genomic sequences between *Fat1* and *F11* genes are complete, and yield no remnants of *Mtnr1a*.

**Figure 3:**
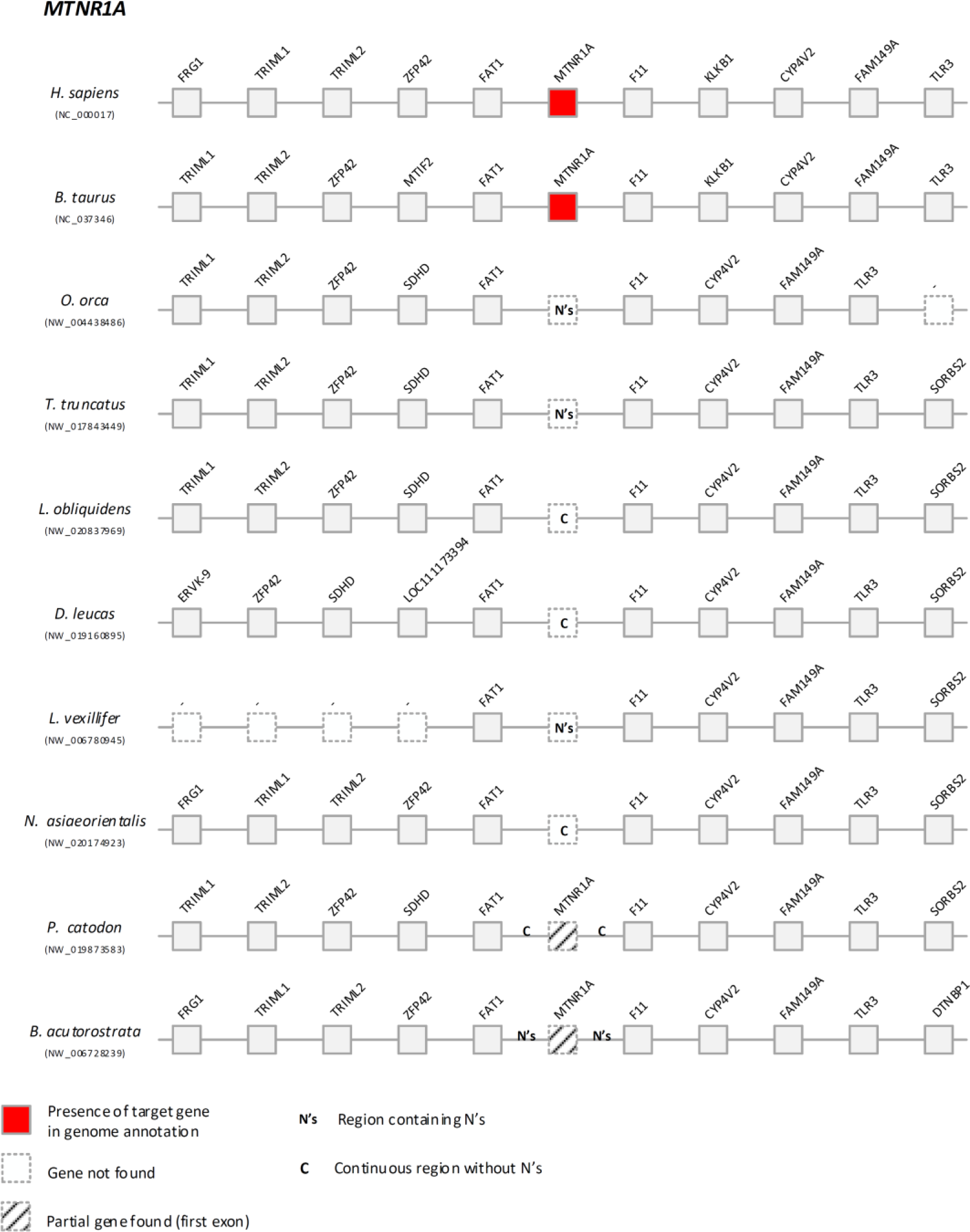
Comparative synteny maps of *Mtnr1a* genomic locus.

In Mysticeti, we were able to deduce the presence of relics of exon 1, yet presenting potentially disruptive mutations (Figure 4). For instance, in *P. catodon*, a 19 nucleotide deletion in exon 1 was found and validated by SRAs (Supplementary Figures 11 and 12). Regarding exon 2, the current assemblies are incomplete and did not allow a decisive conclusion. Yet, overall, the data strongly suggests that *Mtnr1a* is most likely lost or inactivated across Cetacea.

**Figure 4:**
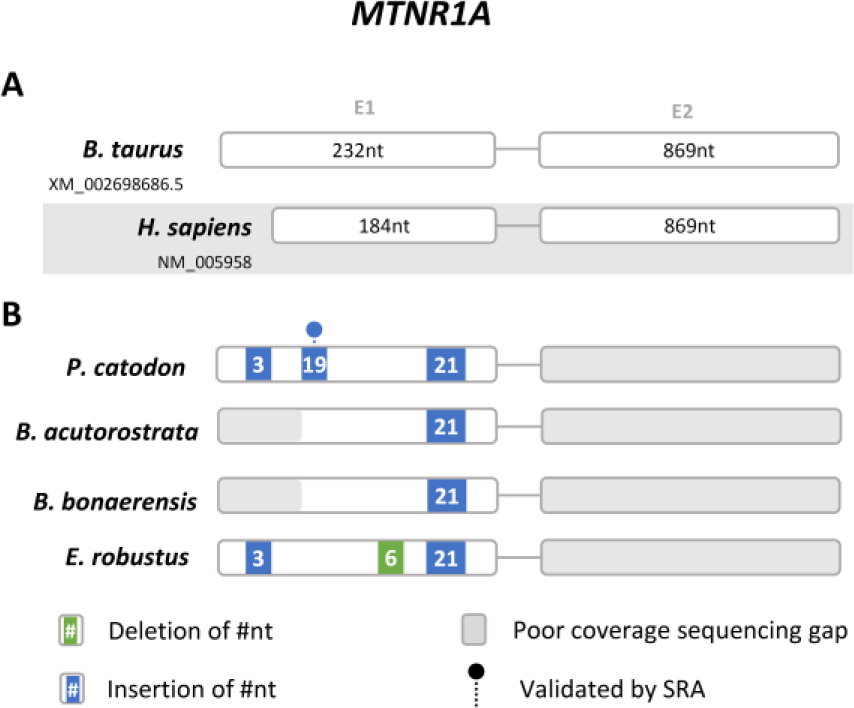
Gene annotation of *Mtnr1a* relics in Mysticeti. **A-** Schematic representation of the gene structure of human and *B. taurus Mtnr1a* gene (not to scale), each box represents an exon and lines represent intronic region. **B-** Schematic representation of the corresponding *Mtnr1a* genes identified in Cetacea and location of the identified insertions and deletions.

In the case of *Mtnr1b*, we retrieved *Mtnr1b*-like sequences in all analysed genomes (Figure 5 and Supplementary Figure 13). Yet, for *L. obliquidens* only the first exon was identified, while no sequencing gaps were detected between the retrieved exon and the neighbouring *Slc36a4* gene (Supplementary Figure 13); exon 2 also remained undetected in *O. orca* and *T. truncatus*, yet the presence of sequencing gaps, between the *Mtnr1b*-like sequence and *Slc36a4*, impaired a final conclusion regarding these species. Further examination of retrieved exon 1 and exon 2 sequences indicated a spectrum of mutations with potential disruptive effects (Figure 5). In exon 1, a conserved stop codon is present in *O. orca* and *S. chinensis*, validated by SRA in the former (Supplementary Figures 14 and 15); a 1 nucleotide insertion in exon 1, generating a premature stop codon, was also found and validated for *B. acutorostrata* and *E. robustus* (Supplementary Figures 14 and 16); and the absence of start codon was detected in *O. orca, T. truncatus, L. obliquidens* and *S. chinensis*, and further validated by SRA in *L. obliquidens* (Supplementary Figures 14 and 17). In exon 2, a trans-species deletion of 1 nucleotide was found in all species with an intact exon, and confirmed by SRA in all species with the exception of *S. chinensis*, shifting the coding frame from this section of the gene (Supplementary Figures 18 and 19). A 282 nucleotide deletion was also identified in *E. robustus* (Supplementary Figure 20).

**Figure 5:**
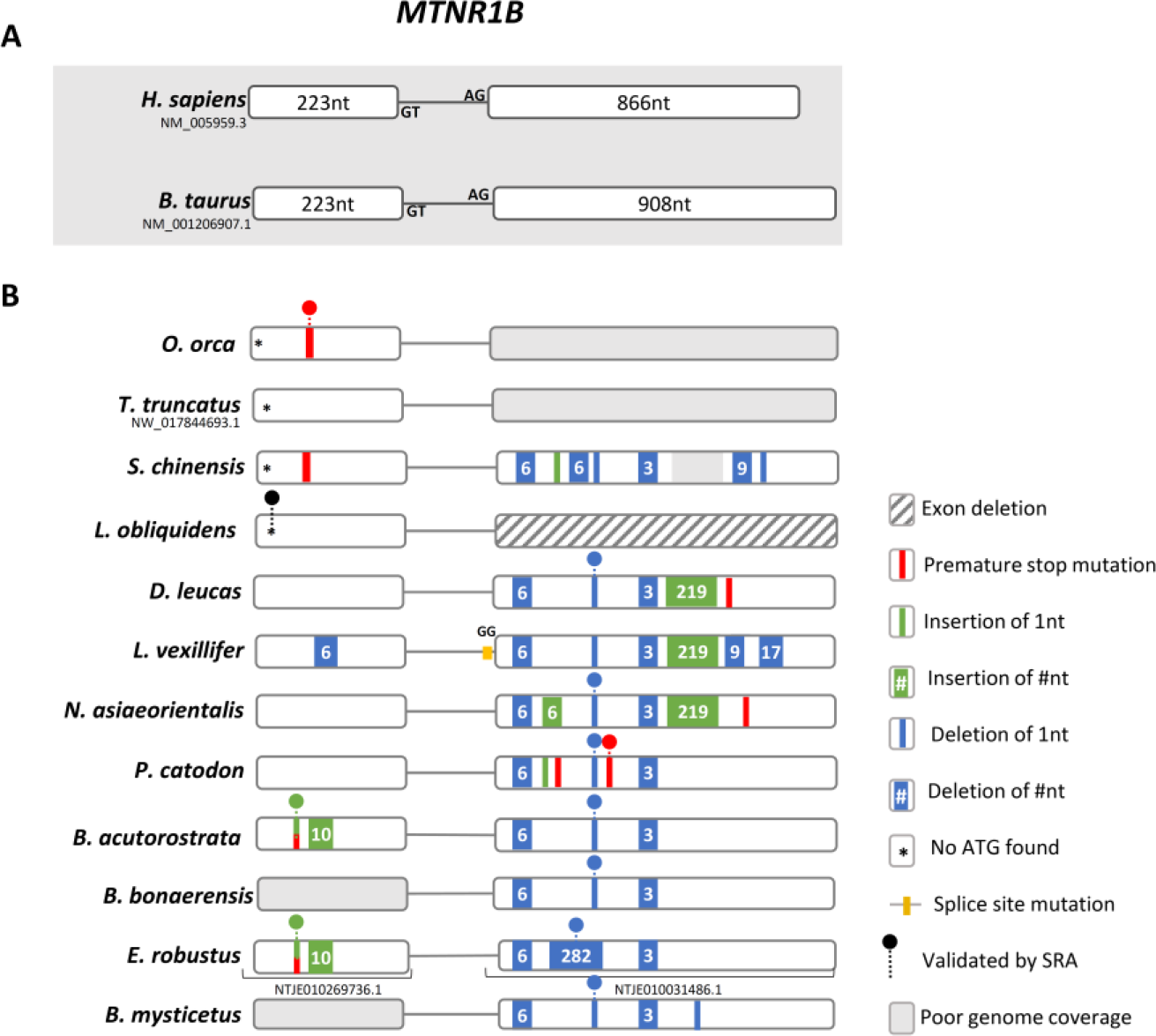
*Mtnr1b* gene annotation. **A-** Schematic representation of the gene structure of human and *B. taurus Mtnr1b* gene (not to scale), each box represents an exon and lines represent intronic region. **B-** Schematic representation of the corresponding *Mtnr1b* genes identified in Cetacea and location of the identified mutations. Non canonical splice sites are indicated above the corresponding annotation.

To further refine our analysis, we next investigated the recently released genome of the *Hippopotamus amphibius* (common hippopotamus)^29^, the closest extant lineage of Cetacea. We were able to deduce the complete ORFs for *Aanat*, *Mtnr1a* and *Mtnr1b* gene orthologues (gene annotations in Supplementary Information). In contrast, in the case of *Asmt* we were able to retrieve partial sequences (exons 2, 5, 7, 8 and 9). Further, searches in SRAs allowed the recovery of exon 1 (Supplementary Figure 21). Regarding *Aanat*, two Aanat-like genes were retrieved for *H. amphibius* as well as *B. taurus* and *Capra aegagrus hircus* (domestic goat), but these are lineage specific duplications (Supplementary Figure 22). Phylogenetic analysis of *Aanat*, *Mtnr1a* and *Mtnr1b* gene sequences revealed that *H. amphibius* orthologues clustered within Artiodactyls (even-toed ungulates), as expected (Supplementary Figure 22 to 24). These findings indicate that the inactivation of melatonin-related pathways most likely took place in the common ancestor of Cetacea, after the divergence of Hippopotamidae and Cetacea.

## Discussion

Taken together, our results strongly suggest that melatonin production and signalling is impaired in Cetacea (Figure 6). In spite of the lower number disrupting mutations found in *Asmt*, we still find indications of pseudogenization in the analysed Cetacea with the exception of *B. mysticetus*, due to poor genome coverage. Furthermore, we were able to infer that ancestral inactivating mutations (shared between toothed and toothless whales), occurred in *Mtrn1b* and *Aanat* ORFs. These evidences support the anatomical observations that pineal gland is absent or vestigial and that classical melatonin-based circadian rhythmicity is lacking in these species^13,14^. Yet, a paradoxical observation emerging from our findings relates to the reported presence of circulating melatonin in the bottlenose dolphin^14^. Without a functional AANAT or specific receptors it seems unlikely that the reported melatonin levels in the bottlenose dolphin are endogenously and rhythmically produced by the canonical cascade and/or act on MTNR-dependent pathways. Thus, melatonin levels could be explained by alternative metabolic pathways: i.e. an increased activity of other enzyme cascades, including AANAT-independent pathways^4,30^. In effect, an ubiquitous ASMT-like gene has been identified in vertebrates, with an unreported function so far^31,32^. Yet, the remarkable conservation of the putative catalytic domain for SAM binding supports the methyltransferase activity of this ASMT-like enzyme^32^. Additionally, melatonin could be acquired from food sources^4^. In fact, no circadian melatonin fluctuations were indisputably acknowledged for bottlenose dolphin^14^. Thus, other MTNR-independent physiological processes could be at play, such as inflammatory and immune responses suggested to be modulated by melatonin-dependent gene-expression^4^. Also, one cannot disregard the ancient antioxidant capacities of melatonin^4^. In fact, cetaceans have evolved strategies to offset hypoxia induced oxidative stress, mirrored by the mutational tinkering observed in mink whalexs’s genes participating in antioxidant systems (e.g. glutathione metabolism^33^).

**Figure 6:**
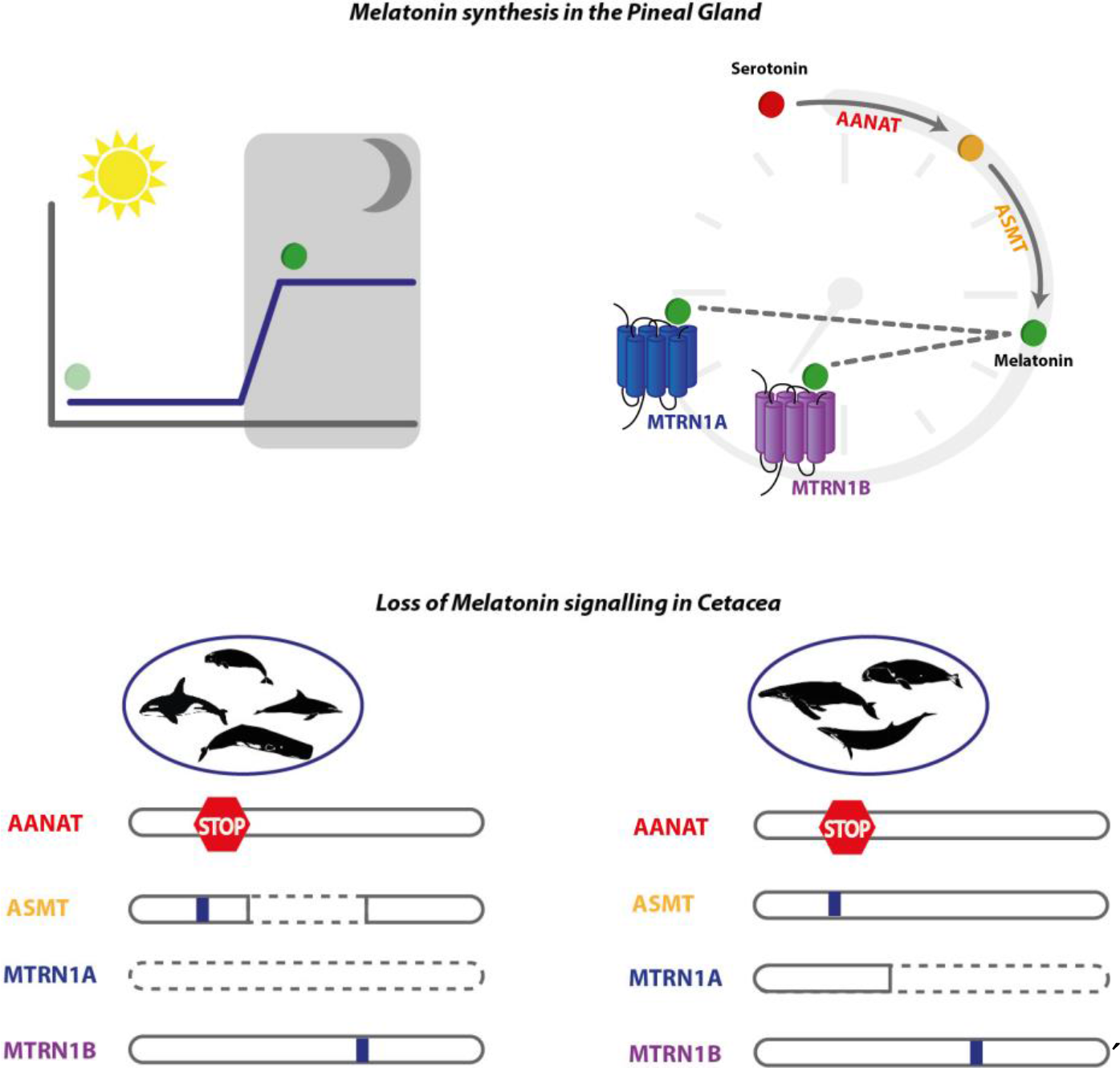
Melatonin synthesis and signalling modules are eroded in Cetacea. Upper Panel: Melatonin synthesis exhibits daily rhythmicity with a production peak at night (left); AANAT and ASMT sequentially convert serotonin into melatonin, which serves as ligand to two high affinity G-protein couple receptors, MTNR1A and MTNR1B (right). Lower Panel: Analysed genes from toothed whales (Odontoceti, left) and baleen whales (Mysticeti, right), premature stops, alternative splicing, indels (blue box), and complete gene erosion (dashed line), are depicted.

What would be the trade-off of losing melatonin synthesis and signalling modules? Melatonin is commonly known for participating in the maintenance of circadian rhythms. Yet, in animals with impaired melatonin synthesis, sleep/wake cycles are not fully disrupted^34^. Nonetheless, melatonin does seem to have synchronizing and phase-locking effects, allowing an optimized adjustment to light/dark cycles^34^. Thus, the erosion of the melatonin synthesis and signalling modules could have accompanied cetacean idiosyncratic sleeping and vigilant behaviours^35,36^. On the other hand, this loss might also reflect other specific life-history traits since cetaceans live in a light-limited environment^37,38^. Overall, our findings provide a unique genomic signature suggesting a behavioural adaptation entailed by a cascade of gene loss events in Cetacea^39^.

## Materials and Methods

### Synteny maps

To perform synteny analyses, the human and *B. taurus loci* of *Aanat, Mtnr1a, Mtnr1b* and *Asmt* genes were used as reference. Importantly, in *B. taurus* two *Aanat* genes were found, and thus we considered as main reference the gene with NM_177509.2 accession number. After collecting the genomic regions of the reference species, several Cetacea genome assemblies and annotations were inspected and scrutinized using the NCBI browser (*H. sapiens* - GCF_000001405.38, *B. taurus* - GCF_002263795.1, *O. orca* - GCF_000331955.1, *B. acutorostrata* - GCF_000493695.1, *D. leucas* - GCF_002288925.1, *T. truncatus* - GCF_001922835.1, *P. catodon* - GCF_002837175.1, *L. vexillifer* - GCF_000442215.1, *N. asiaeorientalis* - GCF_003031525.1). To perform the synteny maps the following procedure was used: 1) five flanking genes from each side of the target gene were collected, 2) only genes neighbouring genes classified as coding were considered (Gene Type: Protein Coding), 3) if the target gene was not found, the genomic region between the direct neighbouring genes (first gene at left and right side) was inspected. The inspection was done in two ways: firstly, via blast searches to determine the total or partial absence of the gene and after, manually to ensure the continuity or disruption of the genomic region (with presence or absence of N’s).

### Sequence retrieval

Genomic sequence retrieval for gene annotation was performed using two strategies: 1In species with fully annotated genomes (0. *orca*, *T. truncatus*, *D. leucas*, *L. vexillifer*, *N. asiaeorientalis*, *P. catodon*, and *B. acutorostrata*), the genomic *locus* of the target gene was collected directly from NCBI. The genomic sequence was inspected, from the 5′neigboring gene to the 3′neigboring gene of the target gene. For example, in the case of *Aanat* the genomic sequence was collected ranging from *UBE20* to *RHBDF*. 2-For species whose genomes were not annotated, the corresponding genomic sequence of the target gene was retrieved, from full genome assemblies, via blastn searches using human and *B. taurus* sequences as query.

To scrutinize the status of the *Aanat*, *Mtnr1a*, *Mtnr1b* and *Asmt* gene sequences in *Hippopotamus amphibius* we built an in-house blast database with the genome assembly (GCA_002995585.1), directly retrieved from the NCBI database. Next, interrogated the inhouse database using the human and *B. taurus*, to retrieve full or partial gene sequences for *H. amphibius*^40^ (blast-n: -word_size 10, -outfmt 6, -num_threads 50). The output results were manually scrutinized and using the qstart, qend, sstart send and bit score options of outfmt6 format of blast software we identified the scaffolds containing each gene. All scaffolds or sets of scaffolds, with evidences of containing the target genes, were gathered and submitted to AUGUSTUS webserver gene prediction^41^. Finally, the annotation files with the CDS and protein sequence of *Aanat, Mtnr1a* and *Mtnr1b* genes were generated (Supplementary Table 1 and Supplementary Gene Annotations). For the *Asmt* gene only exons 2, 5, 7, 8 and 9 were recovered from the genome assembly. Additional searches in SRAs yielded exon 1 (Supplementary Figure 21).

### Gene annotation and mutational validation

For gene annotation, the collected Cetacea genomic sequences were uploaded into Geneious7.1.9. (https://www.geneious.com/) Annotation was performed using as reference the corresponding *B. taurus* gene sequences, with the exception of *Asmt* gene for which the human *ASMT* gene sequence was used as reference due to the presence of multiple expressed isoforms. For each reference sequence all exons were individualized and mapped to the corresponding genomic region in Cetacea, using the built in map to reference tool. The resulting aligned regions were individually inspected to identify ORF abolishing mutations (frameshift, exon deletion, non-canonical splice site and premature stop codons), and concatenated to obtain the Cetacea predicted sequence. Next, at least one conserved ORF abolishing mutation was further validated by the identification of the same mutation in different genomic projects or samples regarding the same species. For this the predicted exonic sequence for each species containing the selected ORF disrupting mutation was used as query in blastn searches in the corresponding available SRAs and Trace Archive in NCBI (only available for *T. truncatus*). Blast hits were collected and uploaded to Geneious and mapped to the corresponding exon, reads poorly aligned or with identity below 95% were removed and mutational status was confirmed when possible, in at least 2 distinct sequencing projects or individuals from the same species. Validation was performed for all species with the exception of *L. vexillifer* for which only one genomic source resulting from a single genome sequencing project is available in NCBI.

### Phylogenetic analysis

The phylogenetic analyses were done using a broad range of mammalian species and using a set of birds as outgroup. To collect the sequences, blastn searches were done (defaults) in NCBI using as reference the annotated human *MTNR1A* (NM_005958.4), *MTNR1B* (NM_005959.3), *AANAT*(NM_001166579.1) nucleotide sequences^40^. In addition, the previously generated *H. amphibius* sequences were used. After sequence collection, a draft alignment was performed and partial and low-quality sequences were removed with Jalview v2.10.5^42^, a total of 79 *Aanat*, 68 *Mtrn1a* and 62 *Mtnr1b* orthologues were included for further analyses (Accession numbers of all datasets can be retrieved in Supplementary Tables 2 to 4). The orthologous gene sets were aligned independently using MAFFT (setting the parameter --auto mode method selection)^43,44^, being automatically selected the method L-INS-I for all datasets. While the alignment of *Aanat* and *Mtrn1a* genes were directly used for phylogenetic analyses, the *Mtnr1b* orthologue alignment required further pre-processing, with all columns containing gaps stripped and removed from the alignment. The phylogenetic analyses were done with the maximum likelihood method, carried in (http://www.atgc-montpellier.fr/phyml/) website and using the PhyML v3.1 software^45^. To determine the best evolutionary model for each set of sequences, the smart model selection (SMS) option was used^46^: selecting HKY85+G+I model for the *Aanat* dataset, and GTR+G+I model for the *Mtrnr1a* and *Mtnr1b* datasets. In addition, to calculate the branch support for each dataset the aBayes method was selected. Finally, the resulting tree were submitted to Dendroscope software^47^, and the bird clade was used to root.

## Acknowledgments

We acknowledge the various Cetacea genome consortiums for sequencing and assembling the genomes. This work was supported by by Project No. 031342 co-financed by COMPETE 2020, Portugal 2020 and the European Union through the ERDF, and by FCT through national funds

## Supplementary information

**Supplementary Figure 1:**
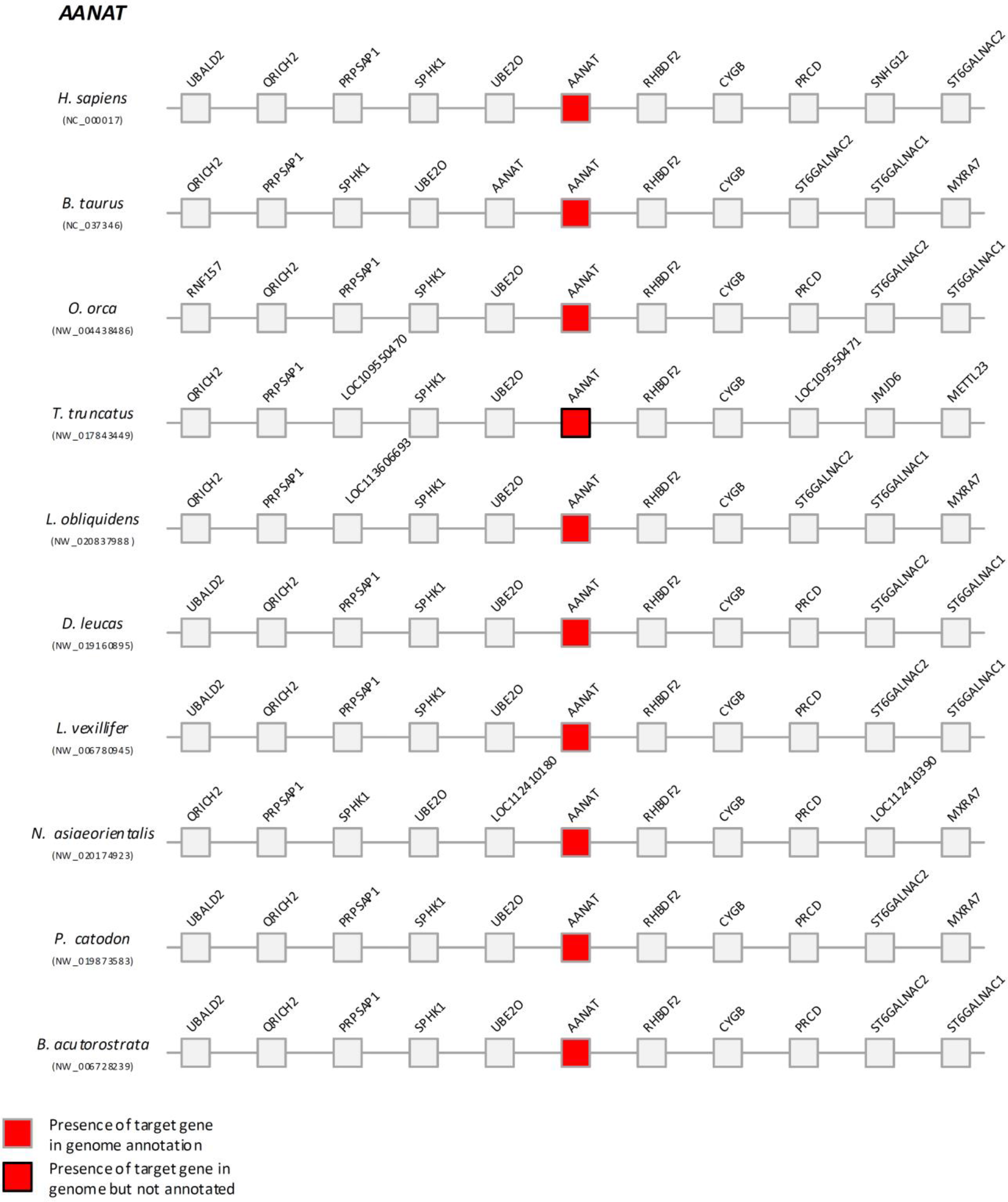
Comparative synteny maps of *Aanat* genomic *locus*.

**Supplementary Figure 2:**
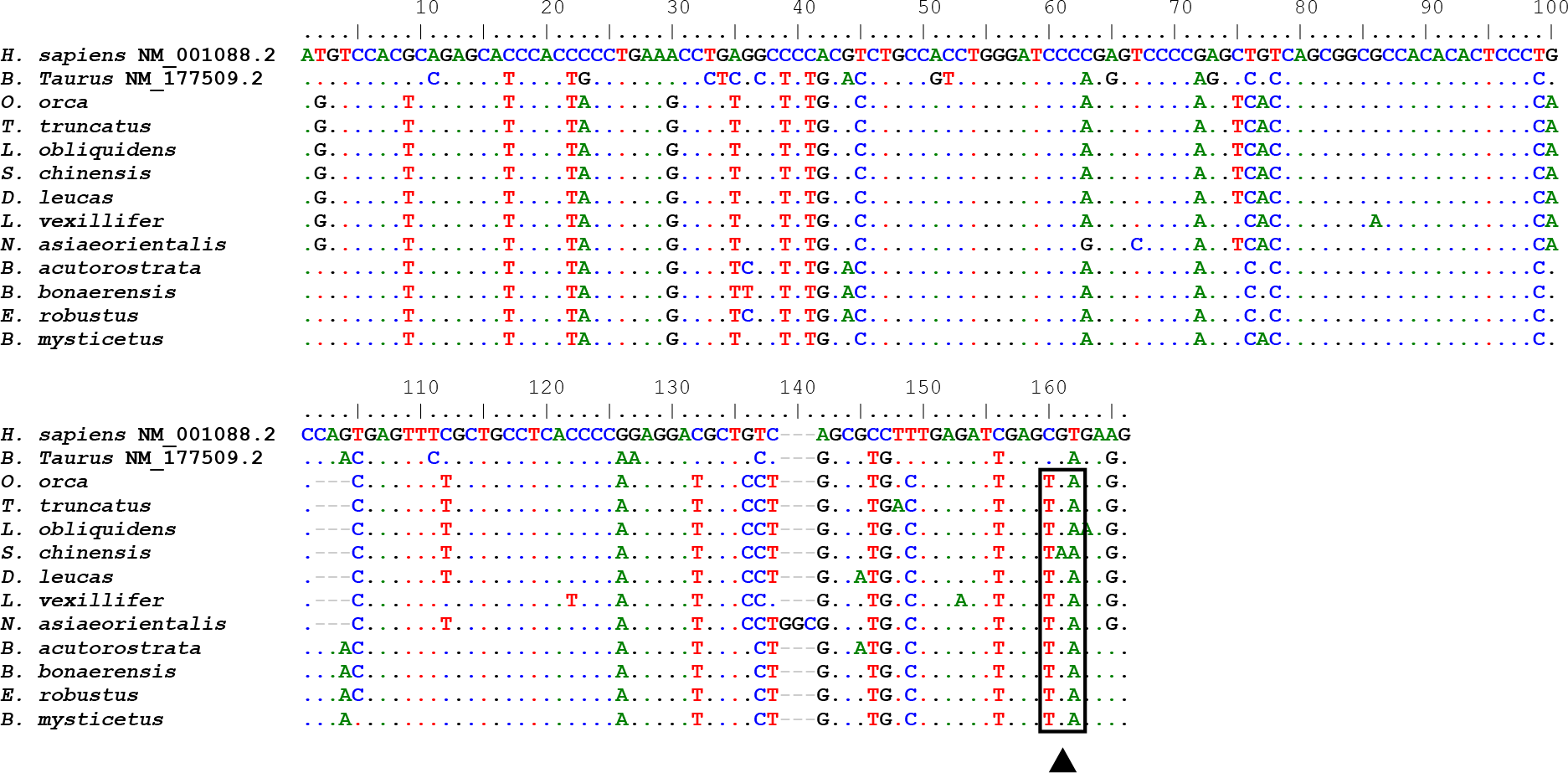
Multiple alignment of the predicted exon 1 of *Aanat* in the listed species. Conserved premature stop mutation, validated with SRA (when available), is represented in the corresponding position with a black arrow.

**Supplementary Figure 3:**
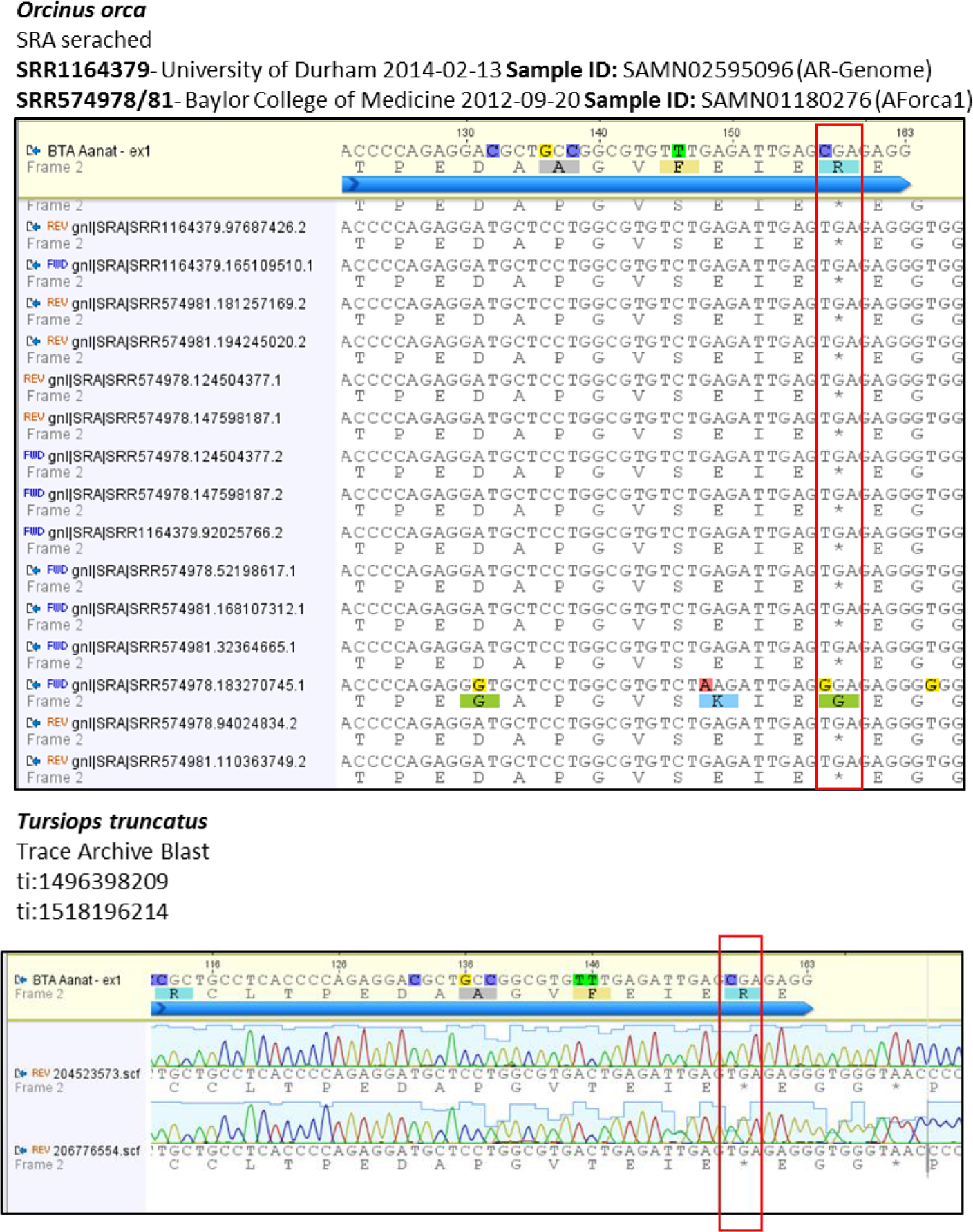
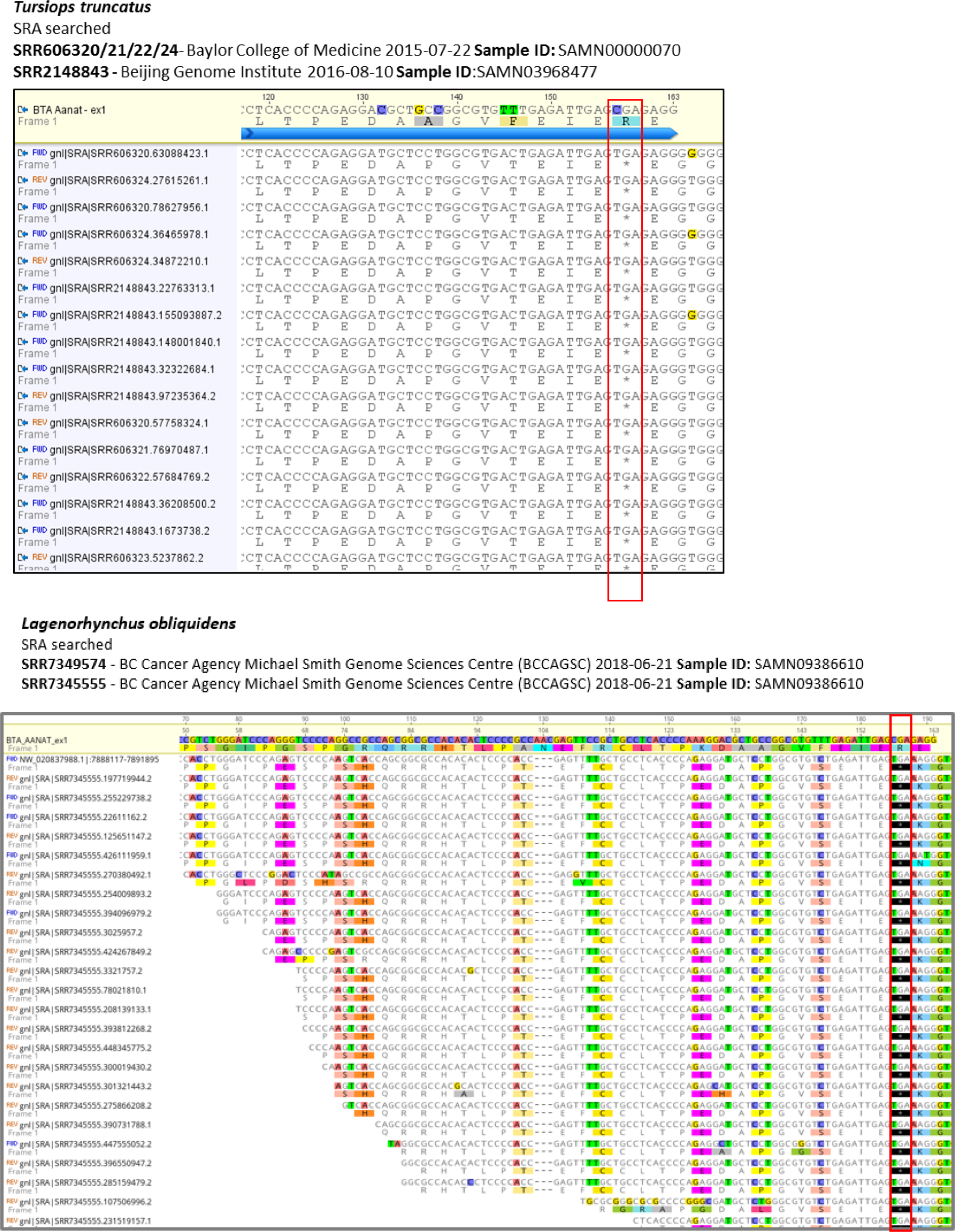
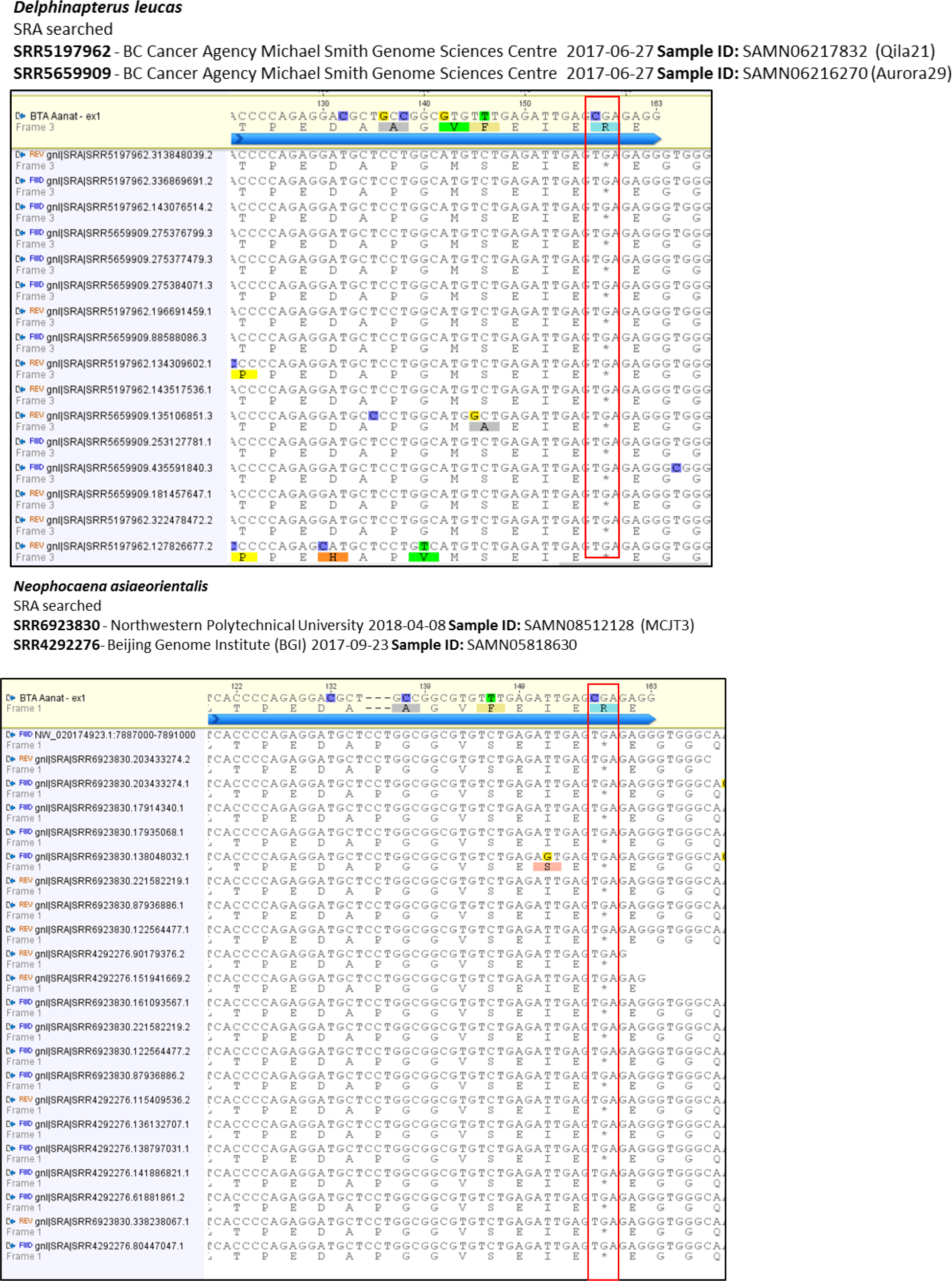
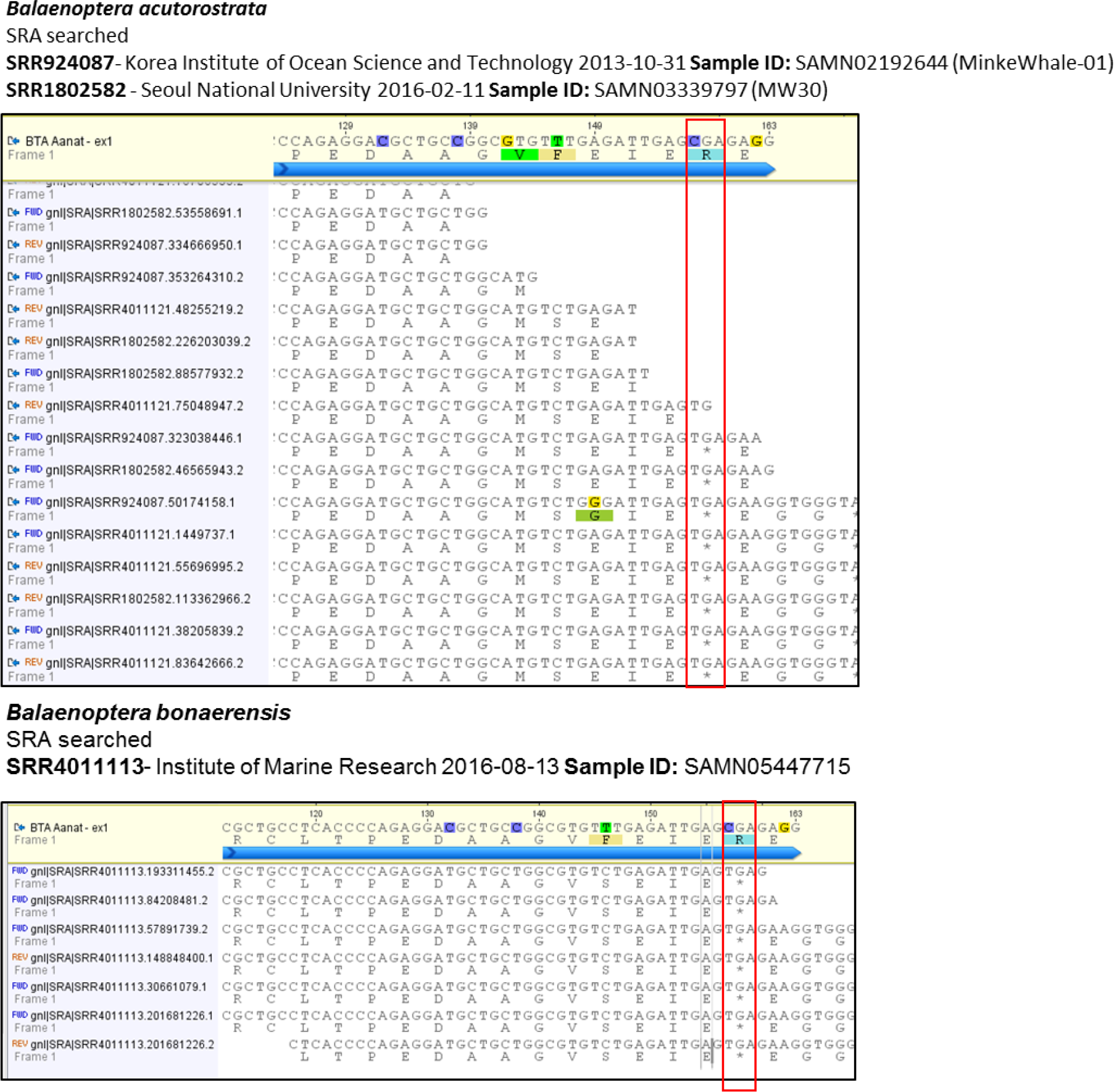
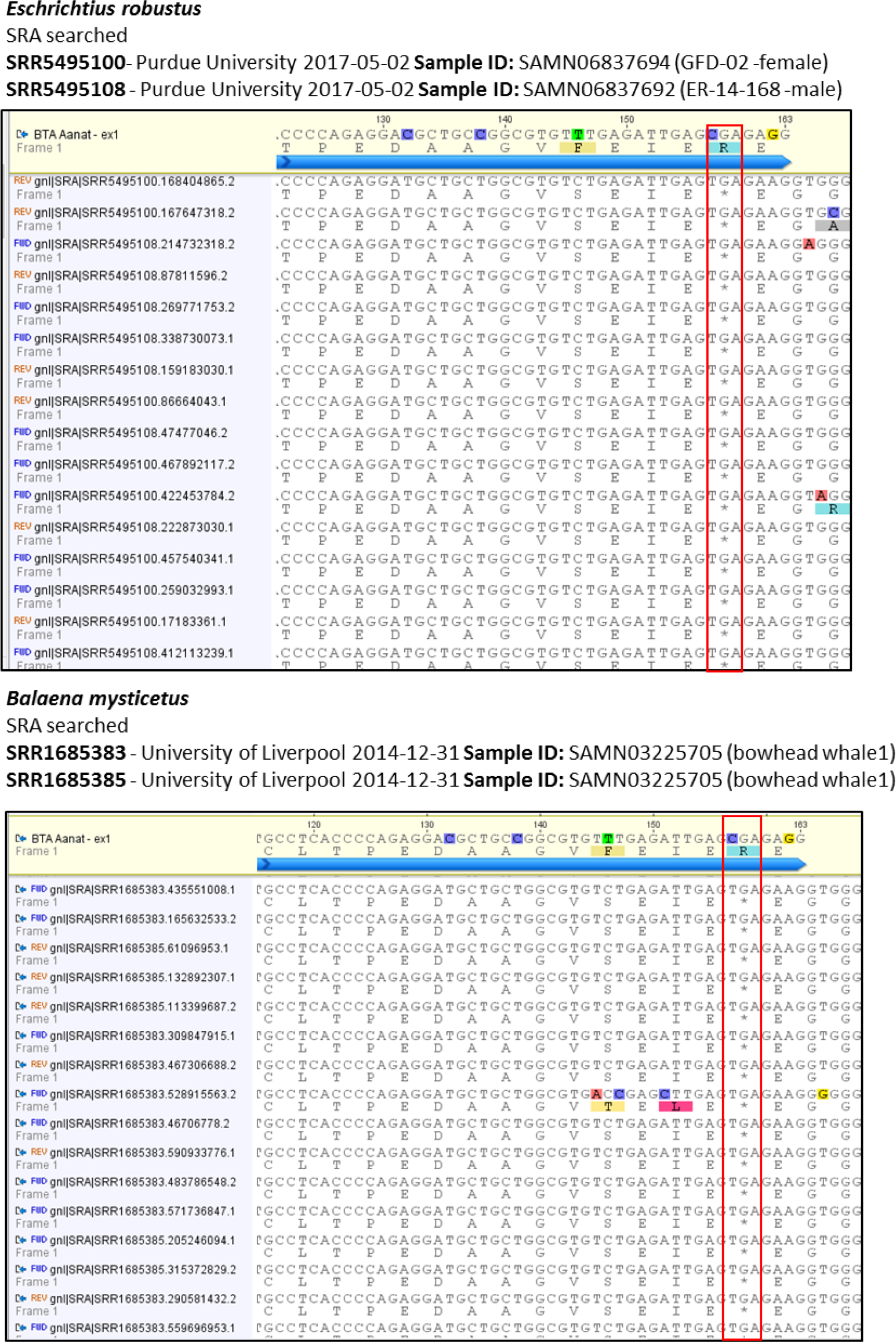
SRA validation for conserved stop mutation (red box) in exon 1 of the *Aanat* gene from *O. orca*, *T. truncatus*, *L. obliquidens*, *D. leucas*, *N. asiaeorientalis*, *B. acutorostrata*, *B. bonaerensis*, *E. robustus* and *B. mysticetus*.

**Supplementary Figure 4:**
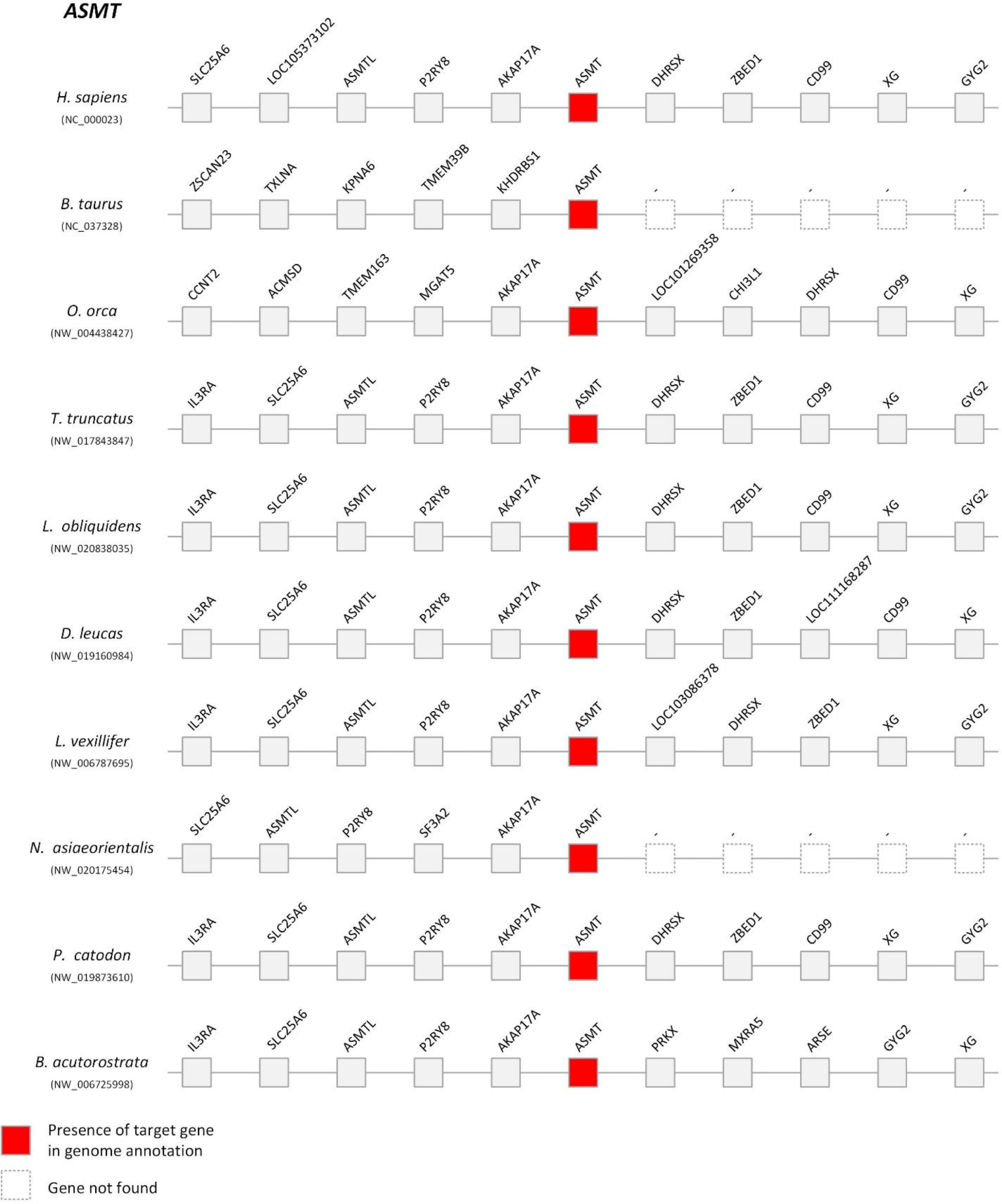
Comparative synteny maps of *Asmt* genomic *locus*.

**Supplementary Figure 5:**
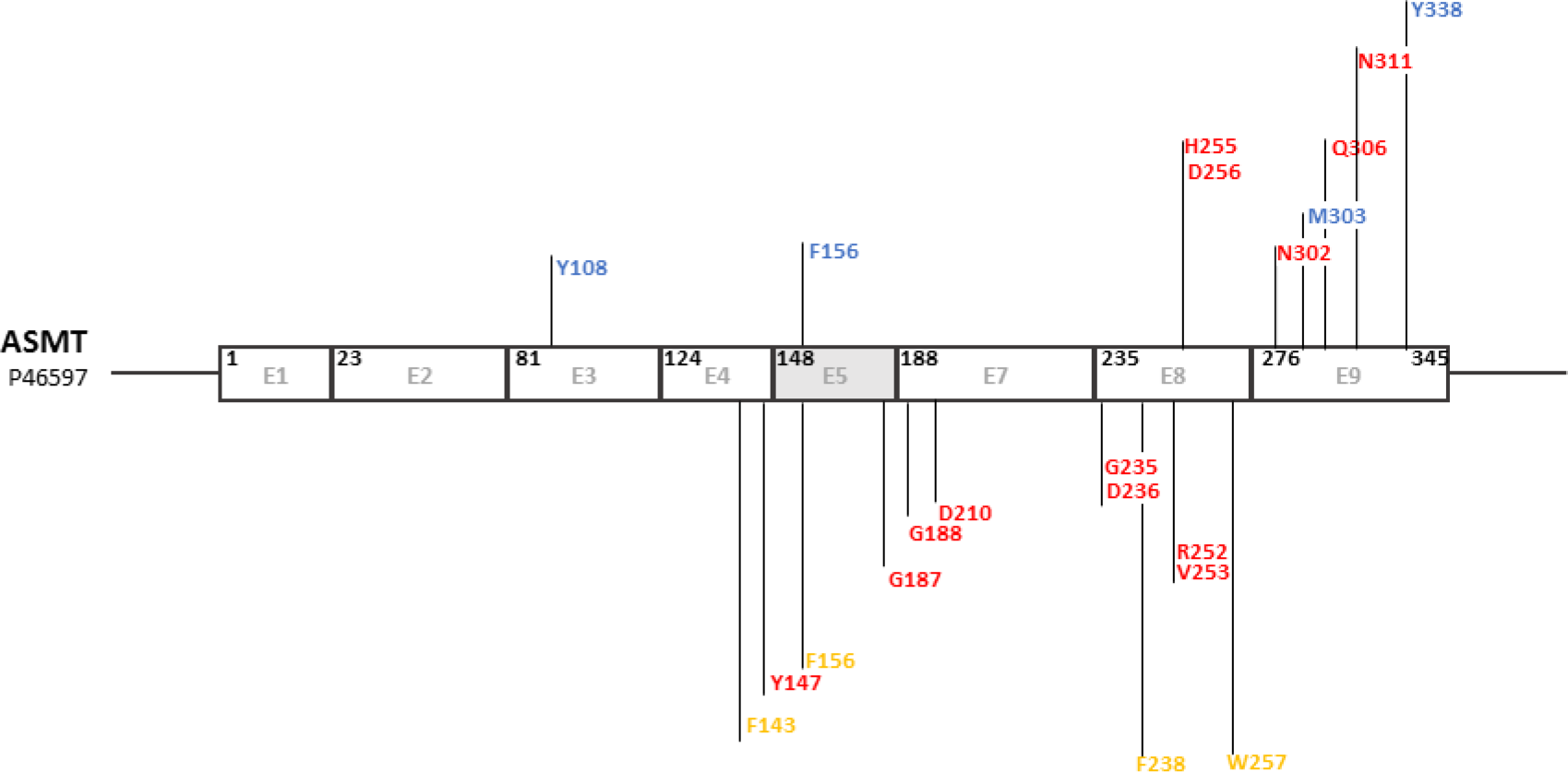
Schematic representation of the Human ASMT isoform 1 (not to scale). On top, in blue, relative location of conserved residues in proximity to the hydroxyl side of *N*-acetyl serotonin (NAS), in red residues that establish H-bonds to NAS. On the bottom in orange conserved aromatic residues that encircle the SAM binding site, in red residues that establish H-Bonds to SAM.

**Supplementary Figure 6:**
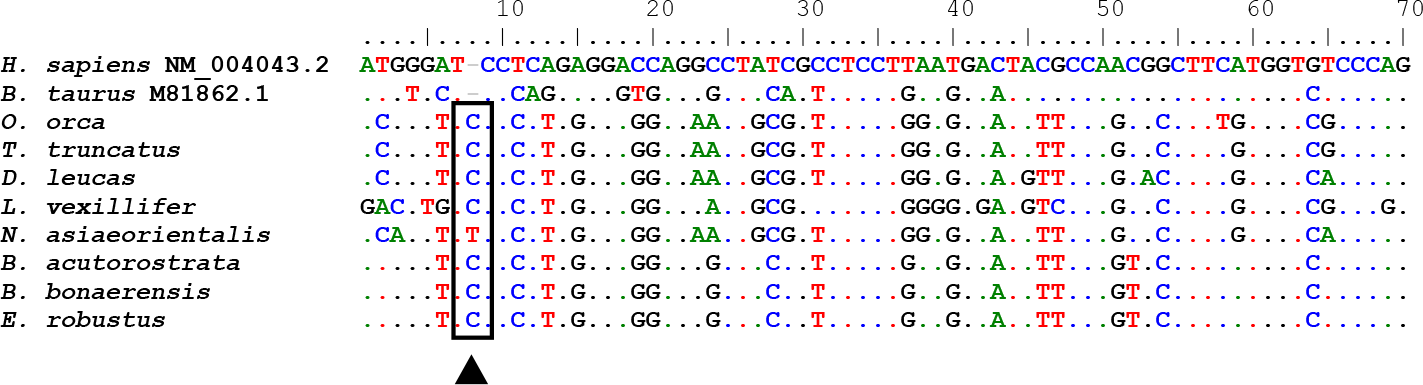
Multiple alignment of the predicted exon 1 of *Asmt* in the listed species. Conserved single nucleotide insertion, validated with SRA (when available), is represented in the corresponding position with a black arrow.

**Supplementary Figure 7:**
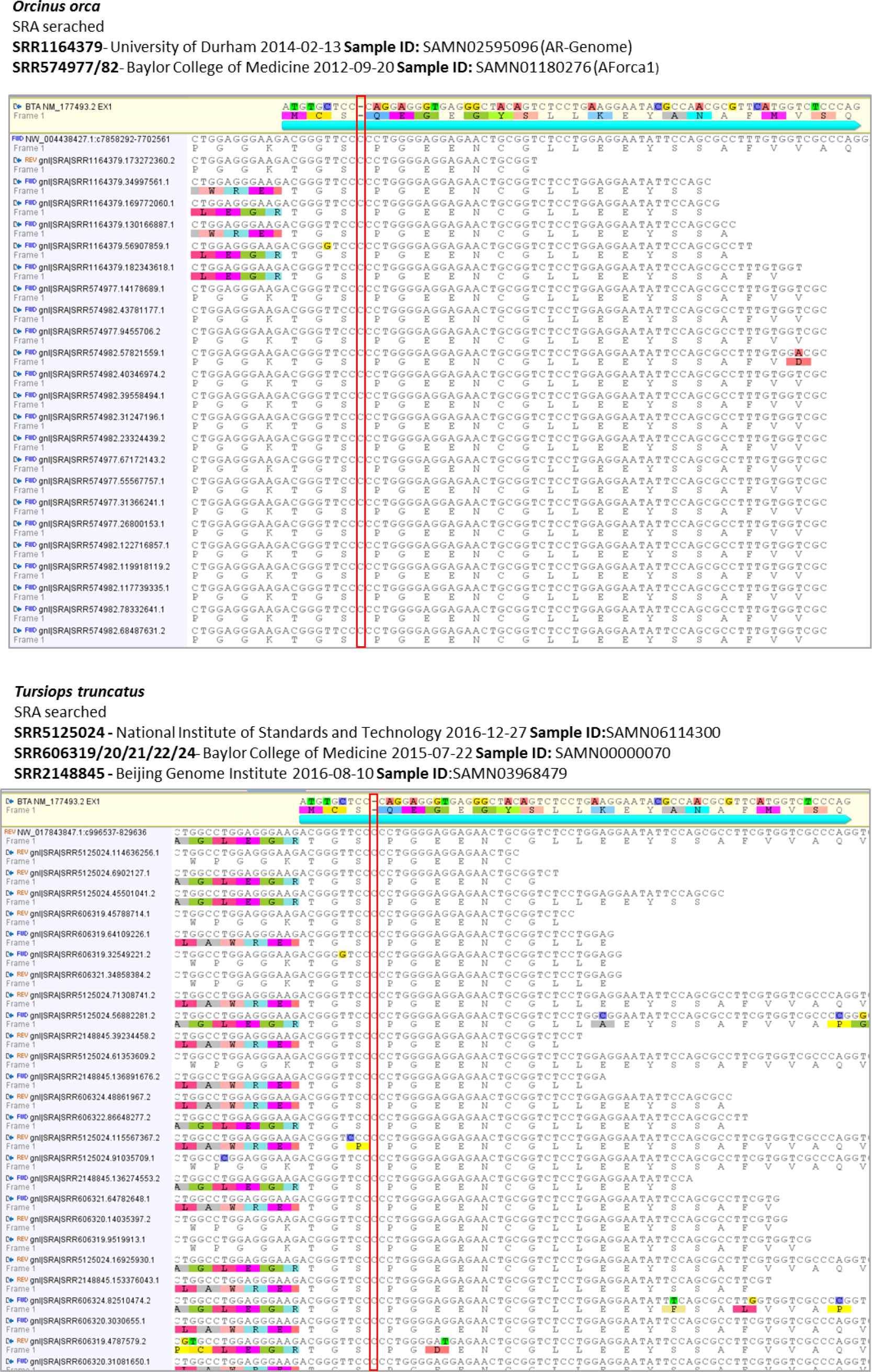
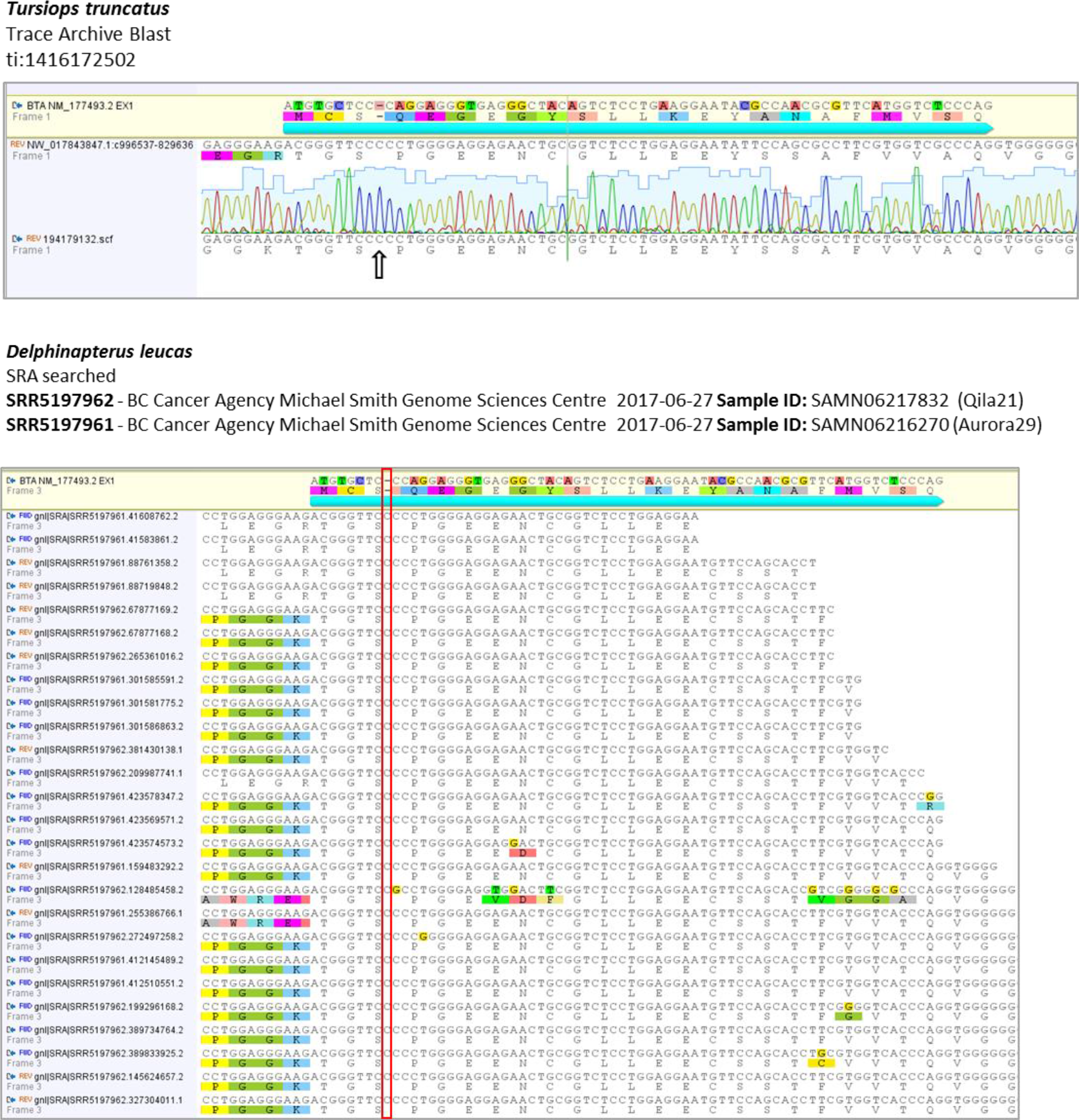
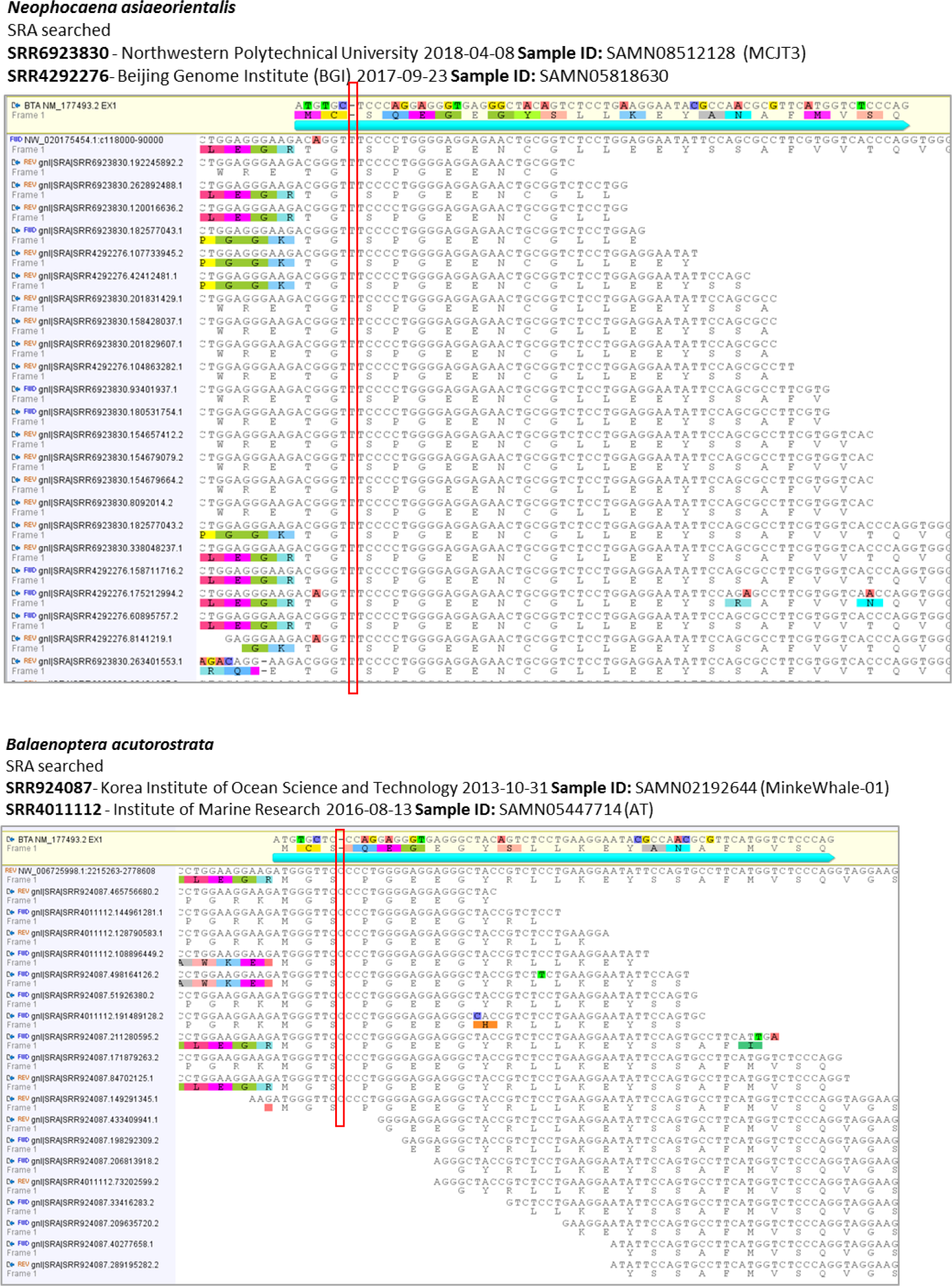
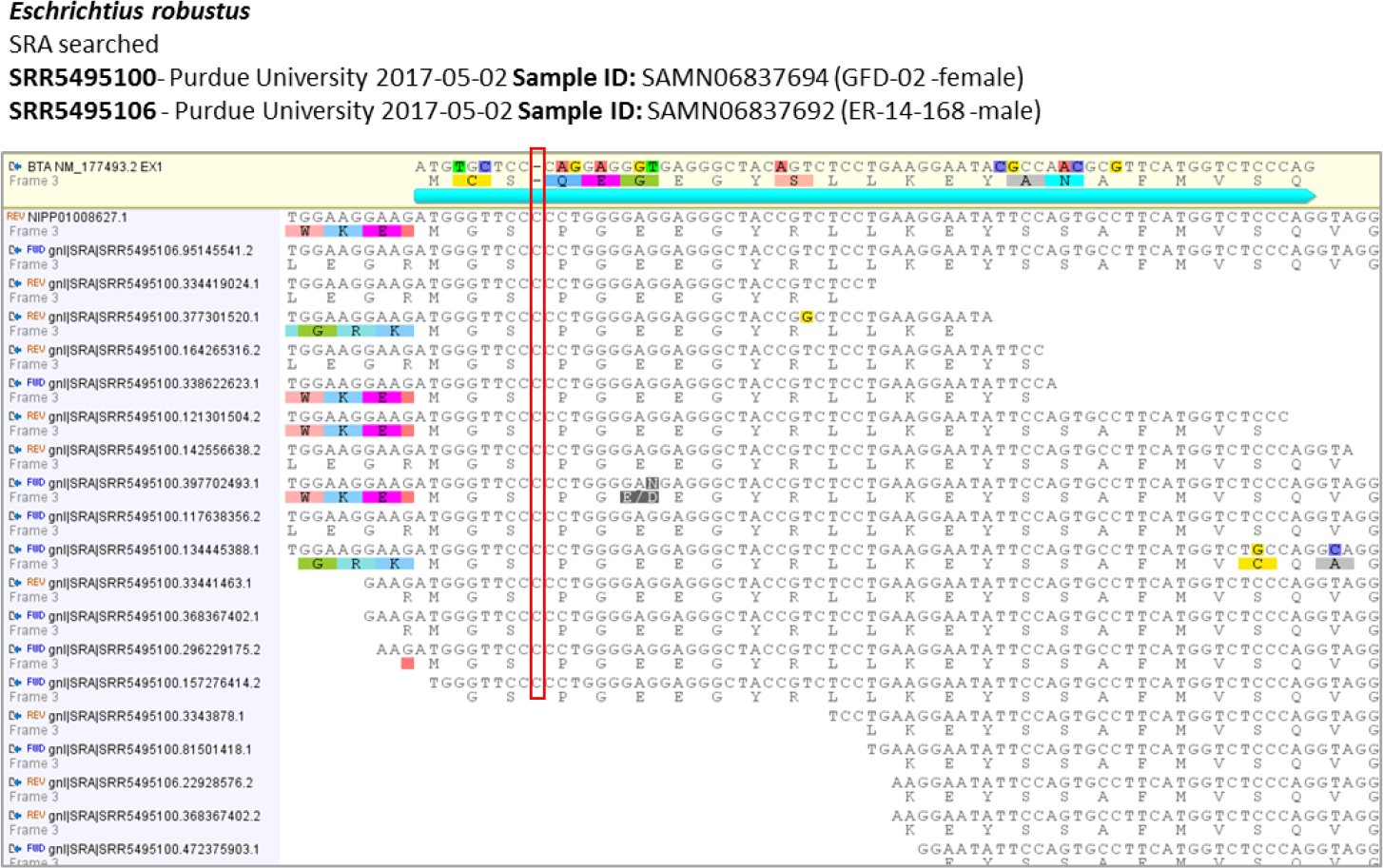
SRA validation for conserved frameshift mutation (red box) in exon 1 of the *Asmt* gene from *O. orca, T. truncatus, D. leucas, N. asiaeorientalis, B. acutorostrata* and *E. robustus*.

**Supplementary Figure 8:**
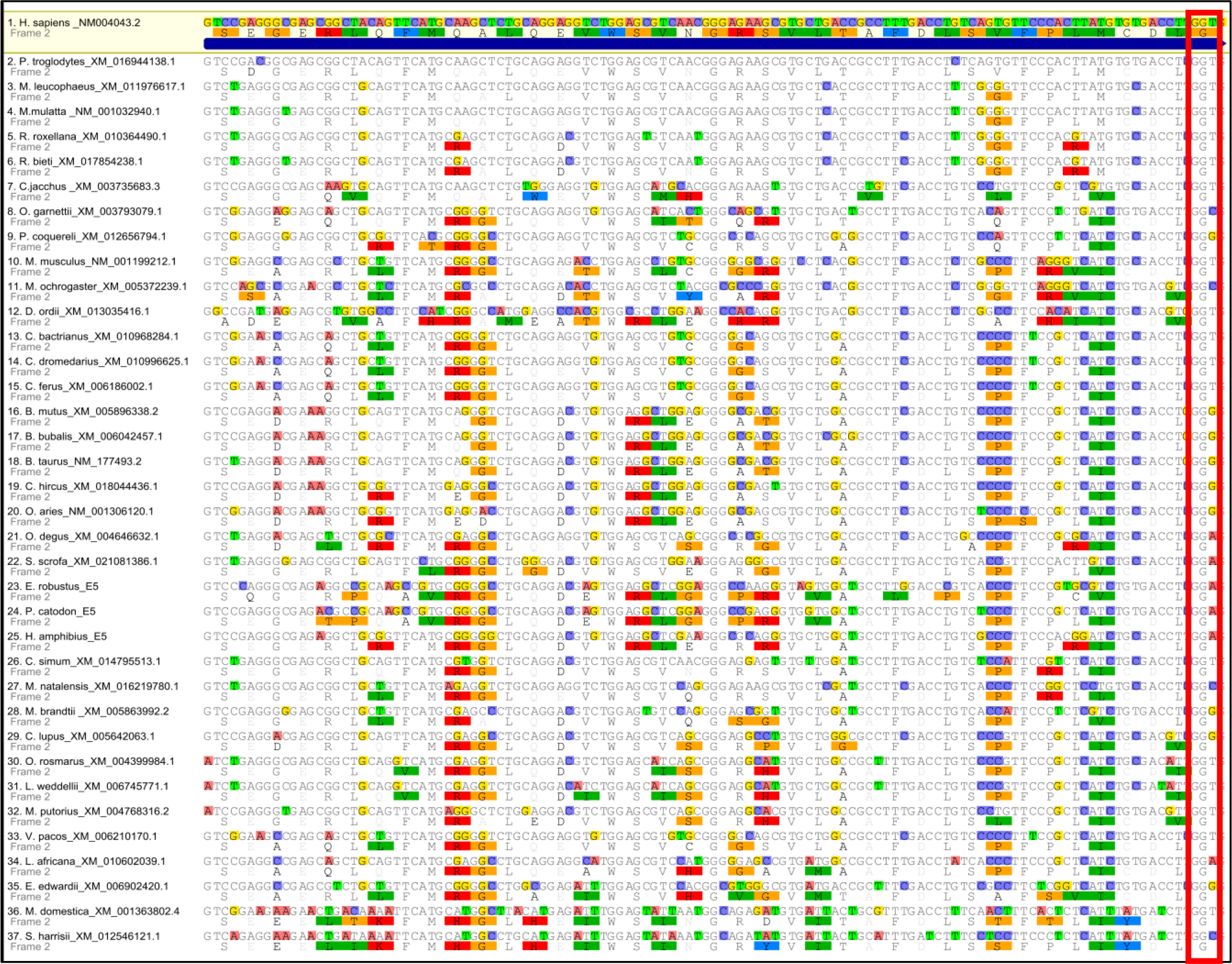
Alignment of exon 5 of *Asmt* in multiple mammals, red box indicates conserved Gly187 participating in the active site of ASMT.

**Supplementary Figure 9:**
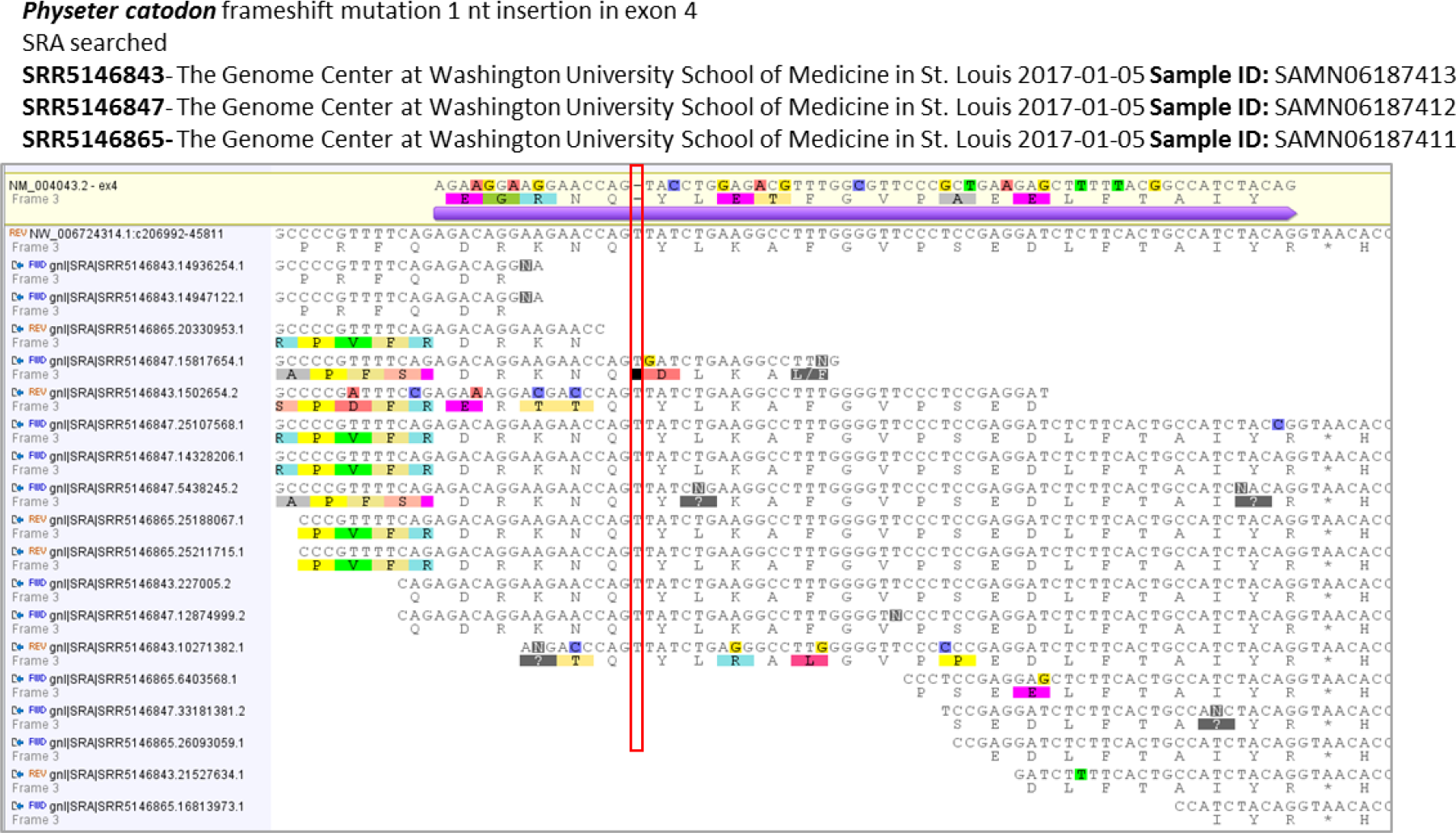
SRA validation for frameshift mutation (red box) in exon 4 of *P. catodon Asmt*.

**Supplementary Figure 10:**
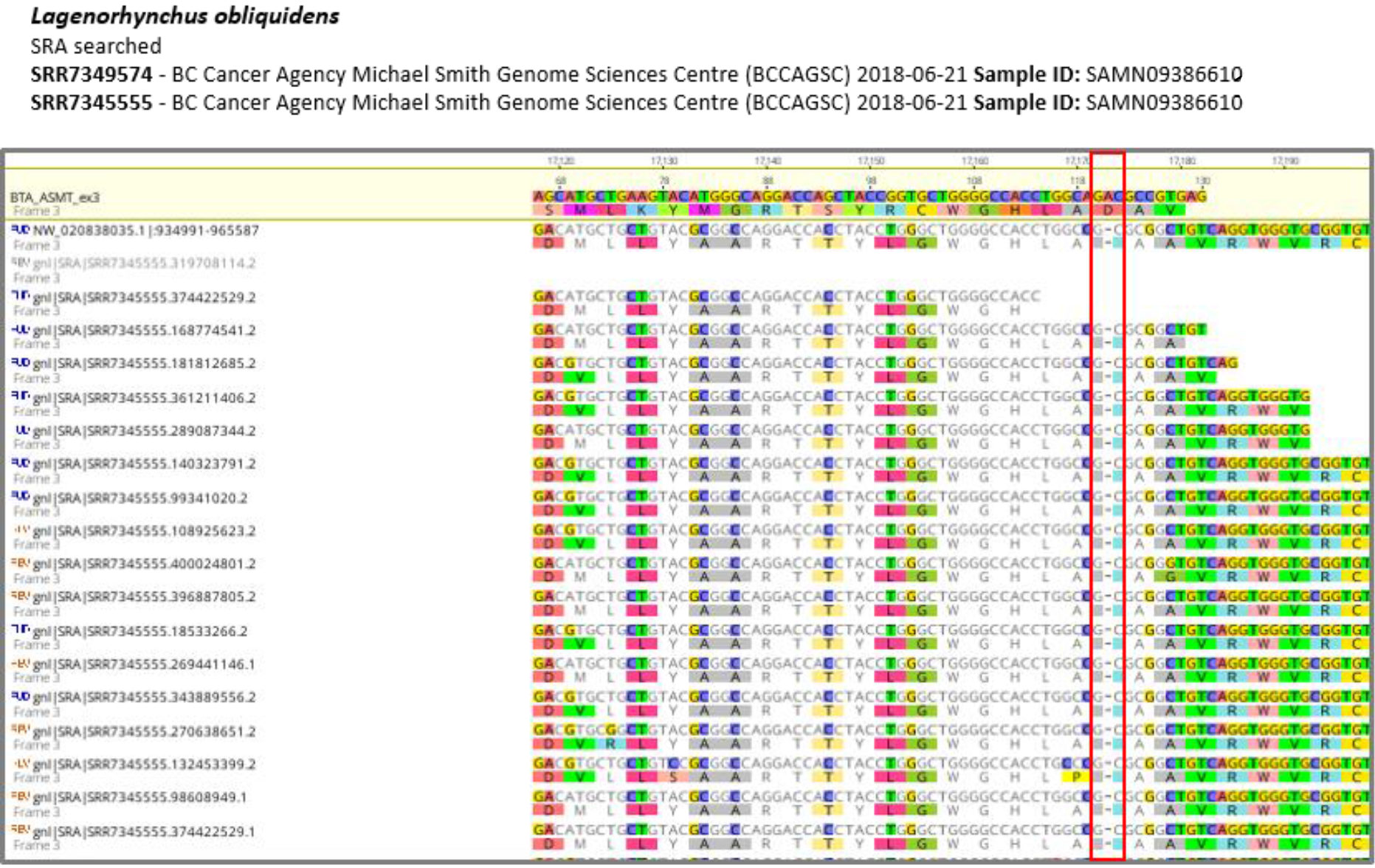
SRA validation for frameshift mutation (red box) in exon 3 of *L*. obliquidens *Asmt*.

**Supplementary Figure 11:**
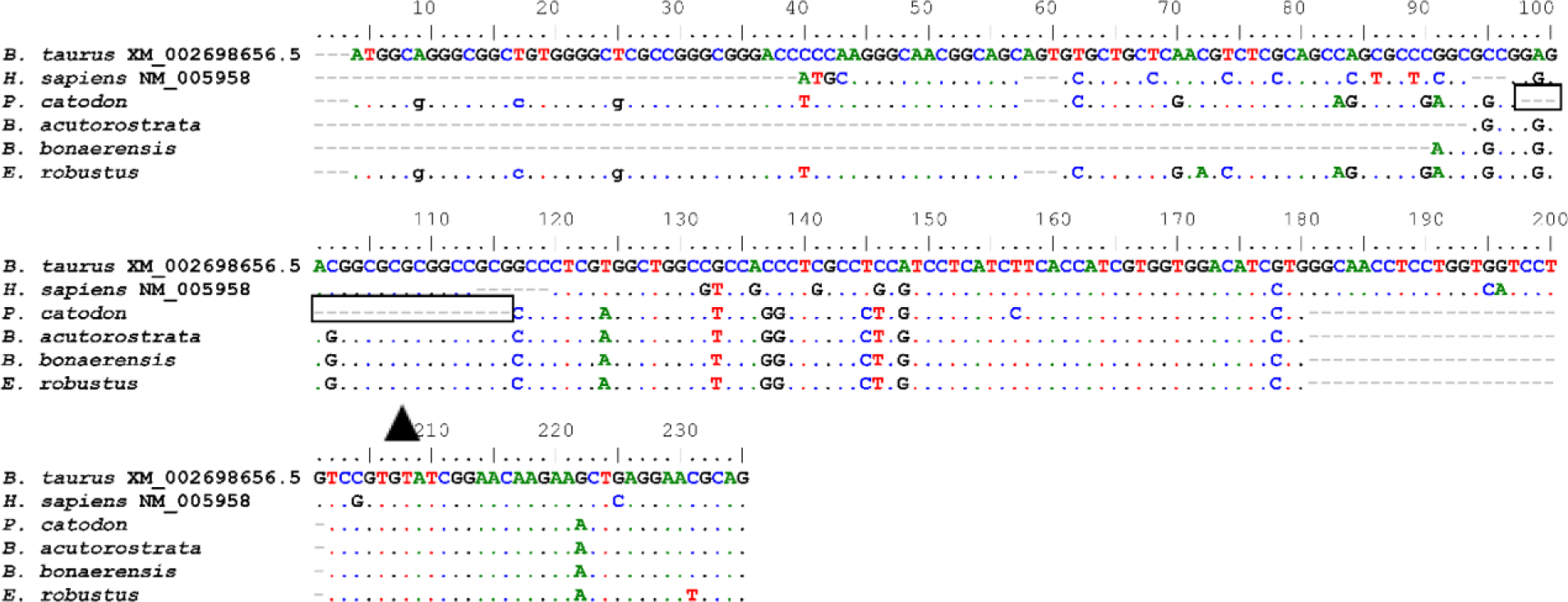
Multiple alignment of the predicted *Mtnr1a* relic sequences (exon 1) in the listed species. Nucleotide insertion of *P. catodon*, validated with SRA, is represented in the corresponding position with a black arrow.

**Supplementary Figure 12:**
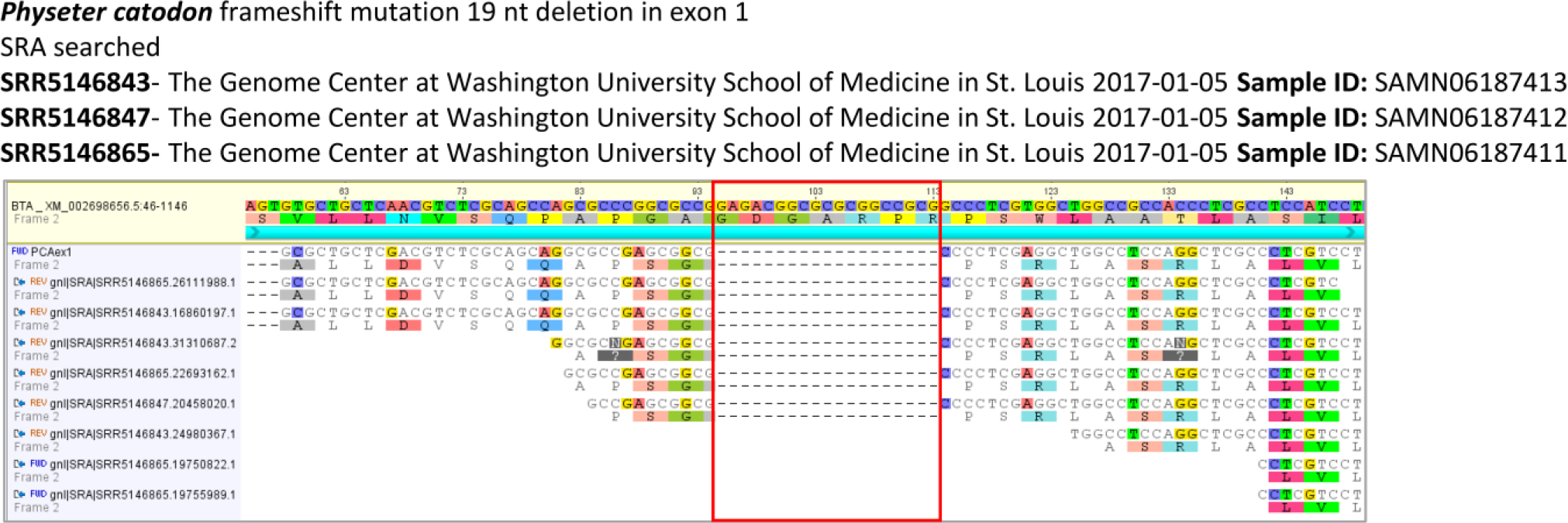
SRA validation for 19 nucleotide deletion in exon 1 of *P. catodon Mtnr1a*.

**Supplementary Figure 13:**
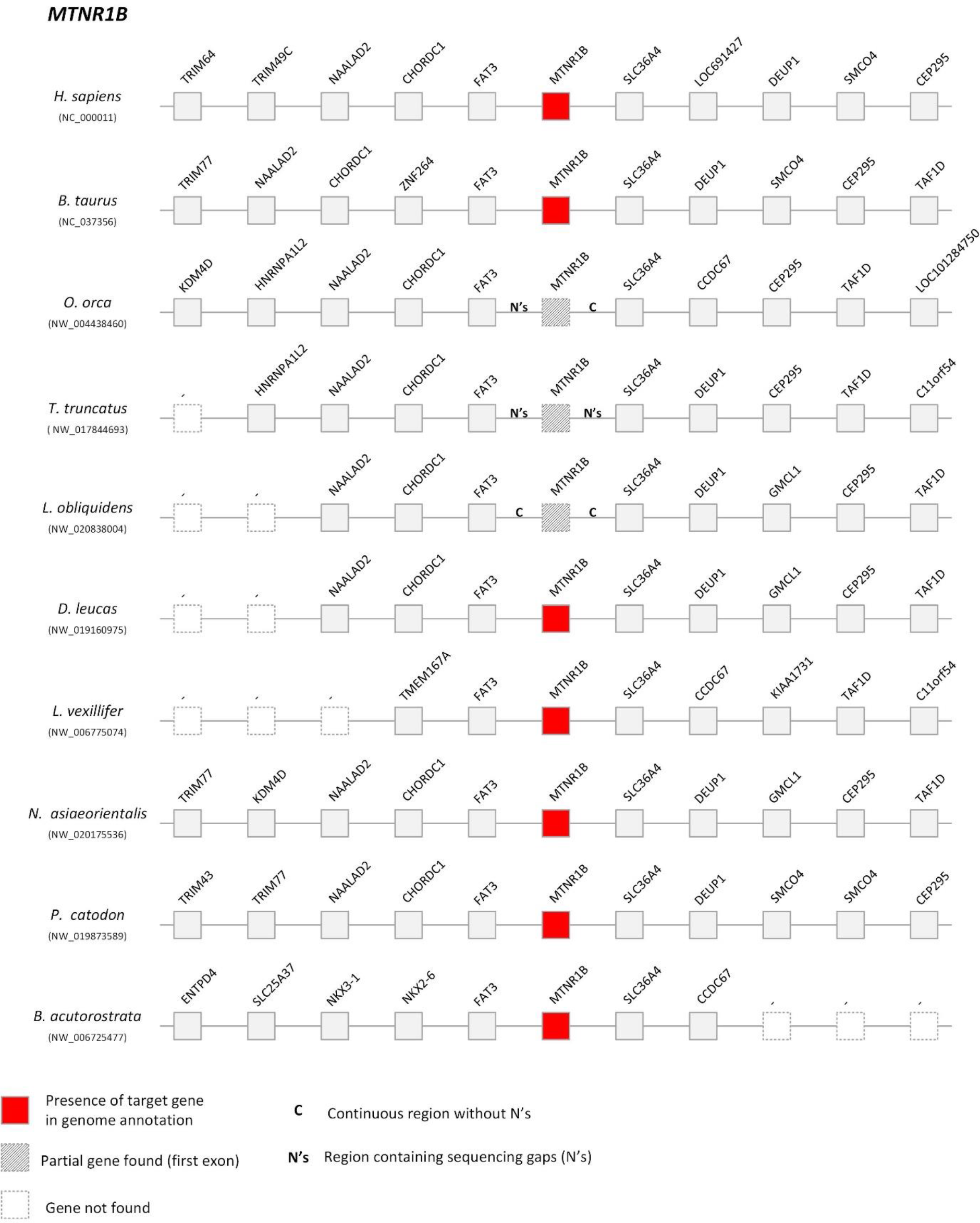
Comparative synteny maps of *Mtnr1b* genomic *locus*.

**Supplementary Figure 14:**
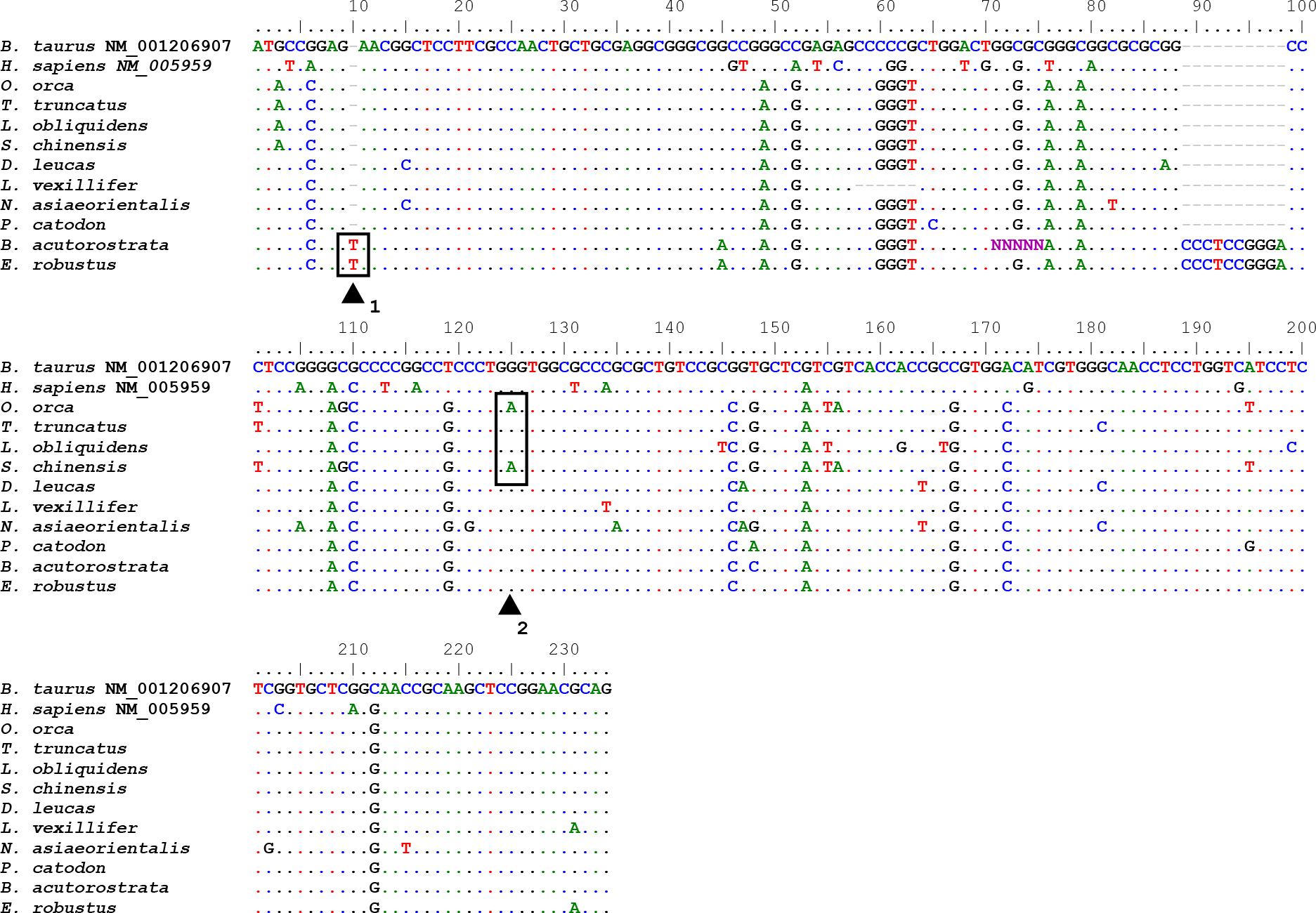
Multiple alignment of the predicted exon 1 of *Mtnr1b* in the listed species. Nucleotide insertion, generating a premature stop codon, in *B. acutorostrata* and *E. robustus* and premature stop codon retrieved in *O. orca* and *S. chinensis*, mutations validated with SRA (when available), are represented in the corresponding position with a black arrow.

**Supplementary Figure 15:**
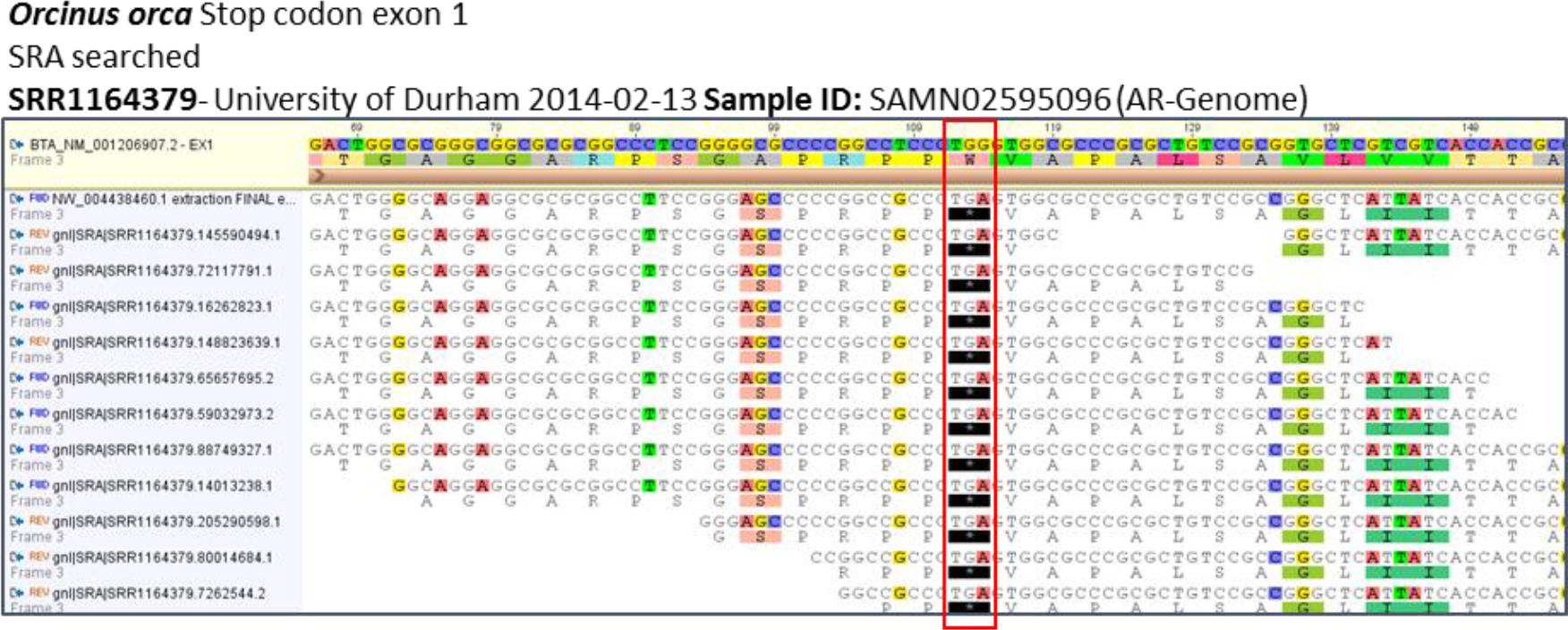
SRA validation for stop mutation (red box) in exon 1 of the *Mntr1b* gene from *O. orca*.

**Supplementary Figure 16:**
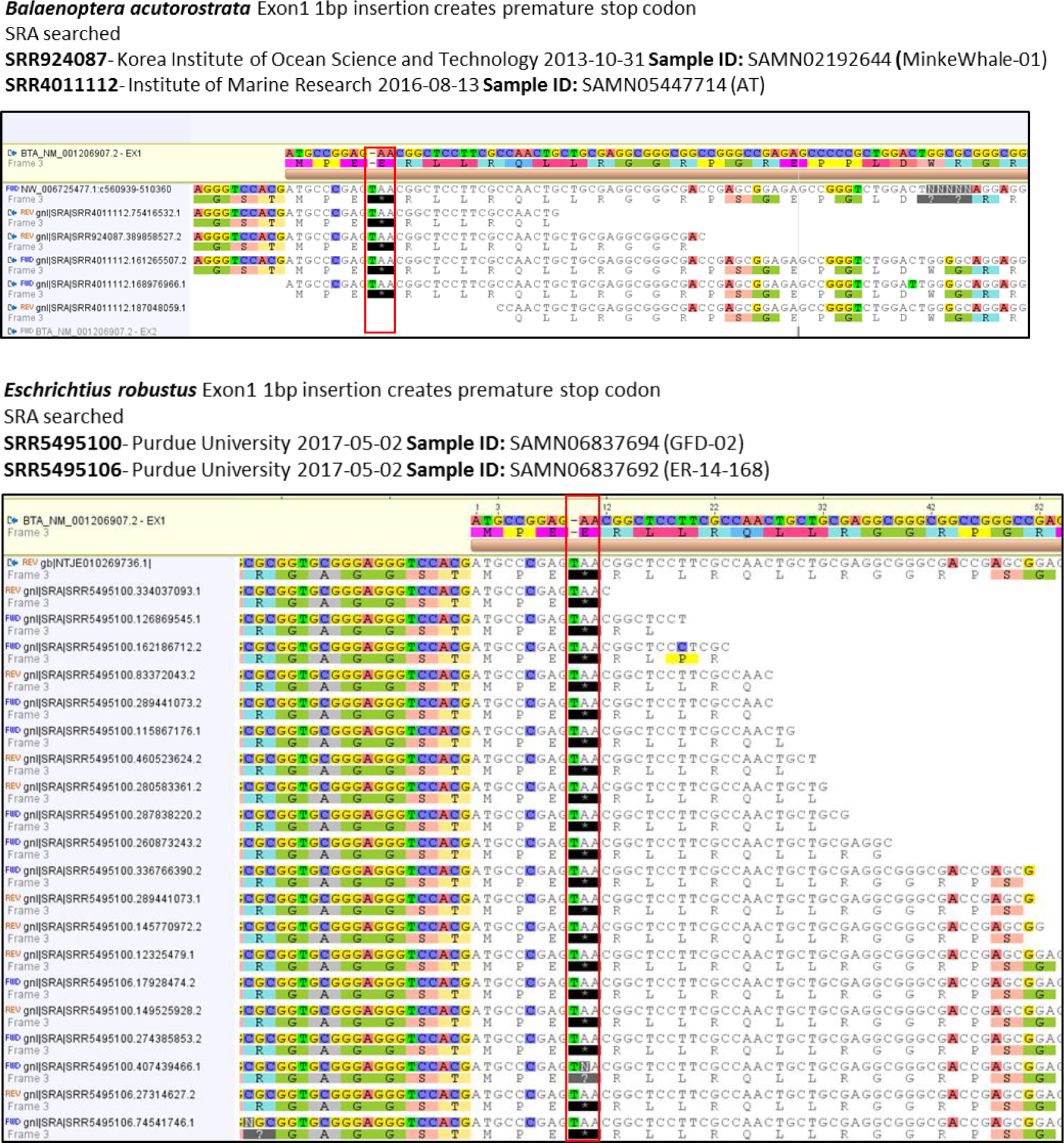
SRA validation for nucleotide insertion generating a premature stop mutation (red box) in exon 1 of the *Mntr1b* gene from *B. acutorostrata* and *E. robustus*.

**Supplementary Figure 17:**
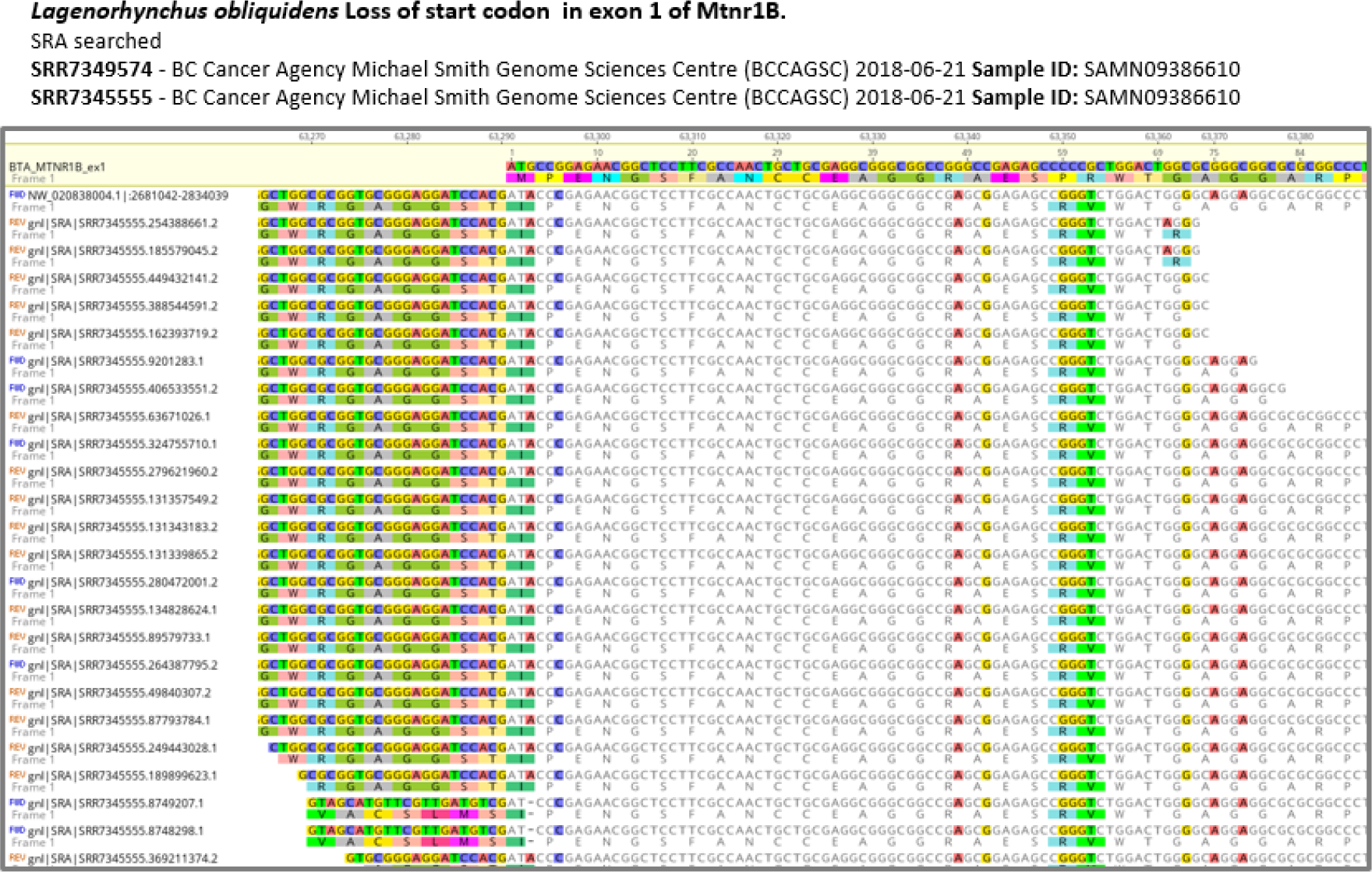
SRA validation for loss of start codon in exon 1 of the *Mntr1b* gene from *L. obliquidens*.

**Supplementary Figure 18:**
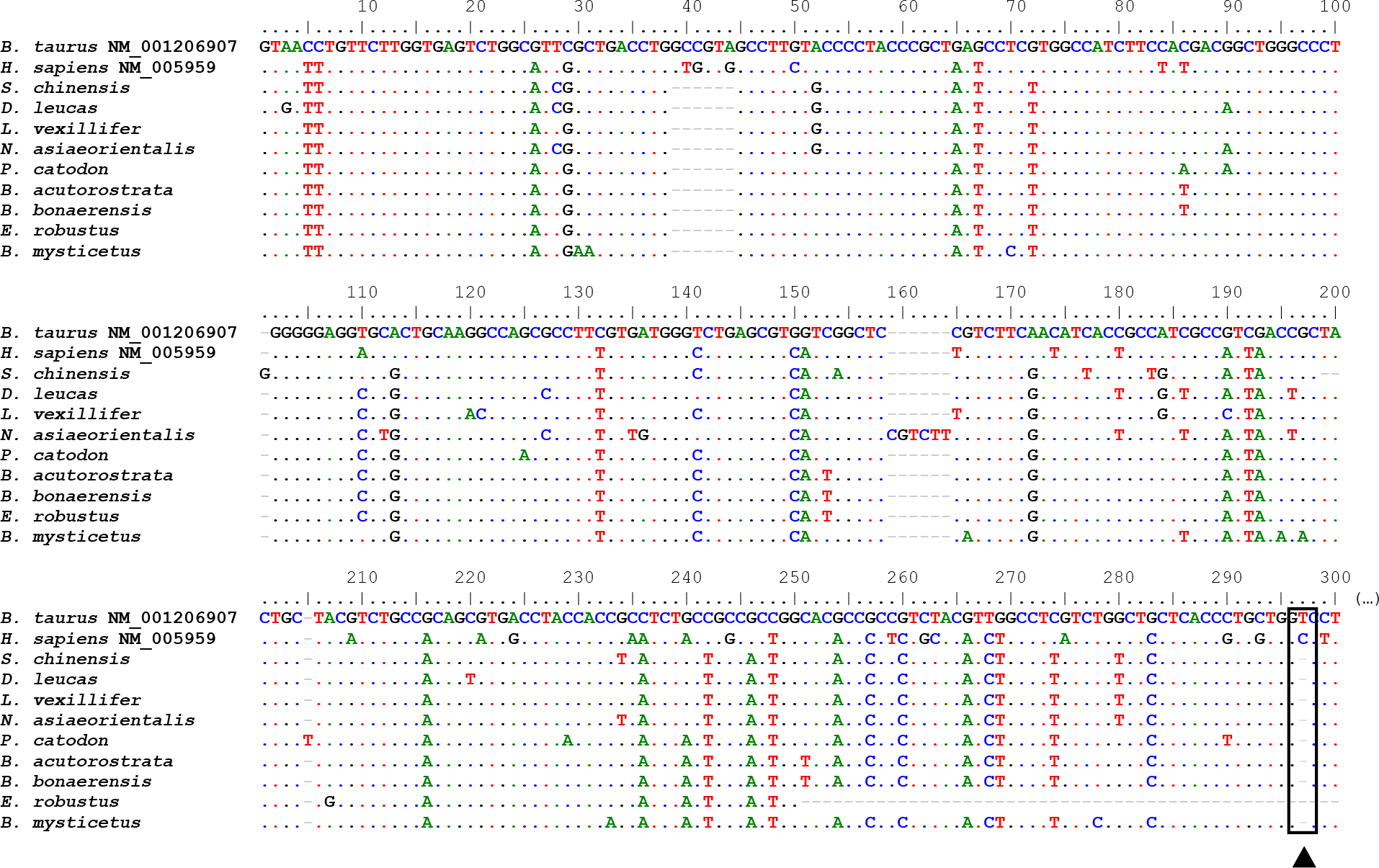
Partial multiple alignment of the predicted exon 2 of *Mtnr1b* in the listed species. Conserved nucleotide deletion, validated with SRA (when available), is represented in the corresponding position with a black arrow.

**Supplementary Figure 19:**
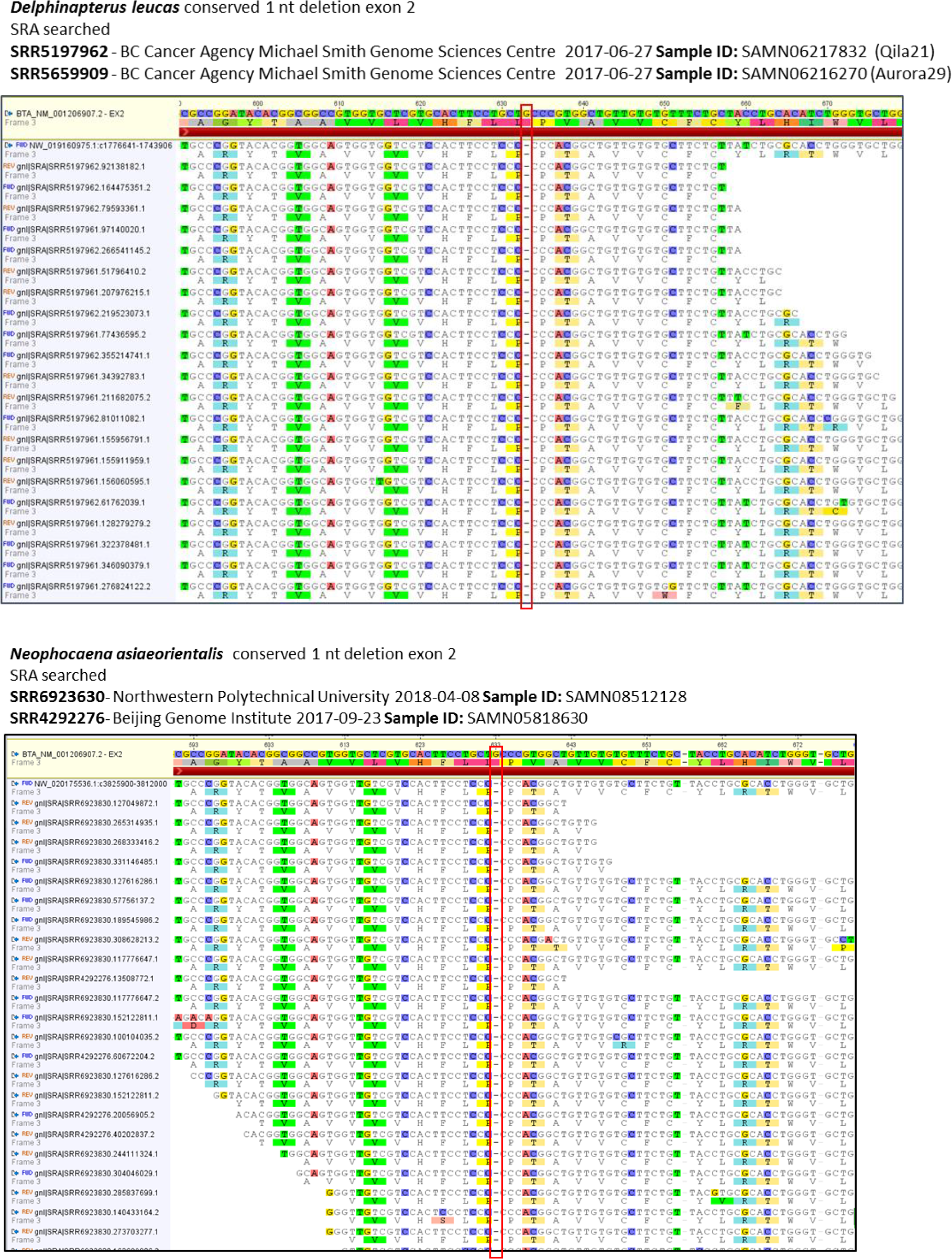
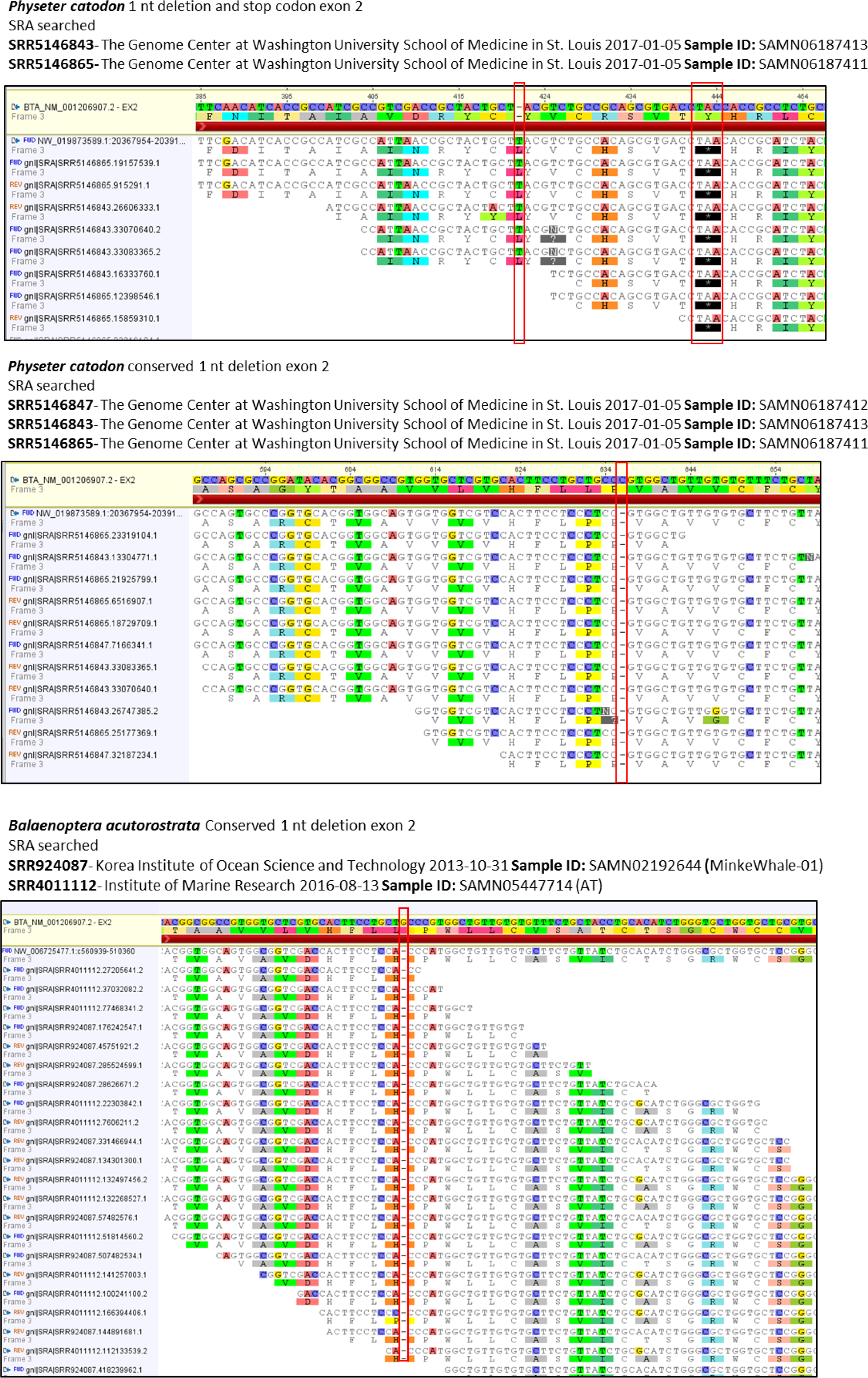
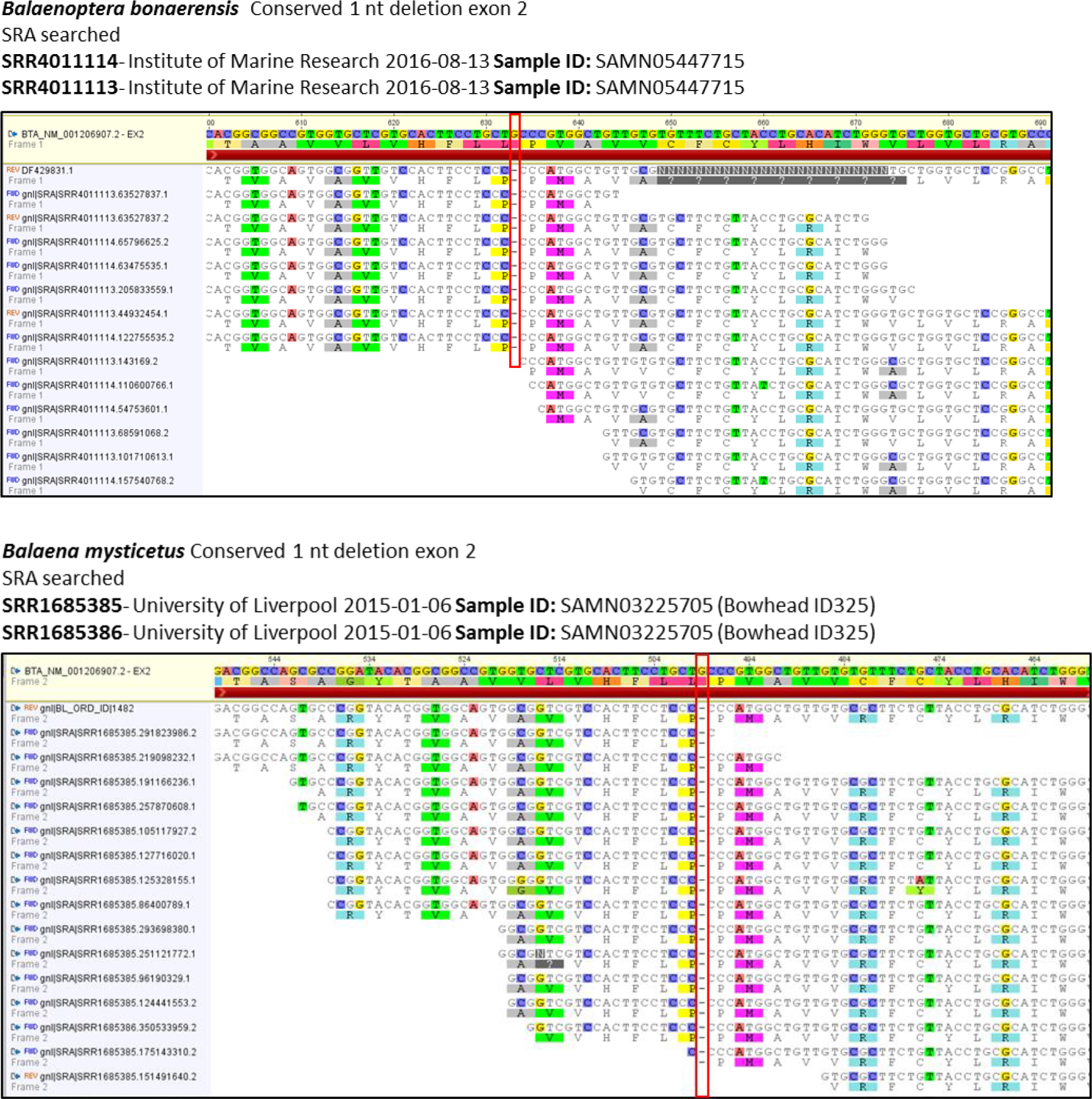
SRA validation for conserved nucleotide deletion in exon 2 of *Mtnr1b* from *D. leucas*, *N. asiaeorientalis*, *P. catodon*, *B. acutorostrata*, *B. bonaerensis* and *B. mysticetus*.

**Supplementary Figure 20:**
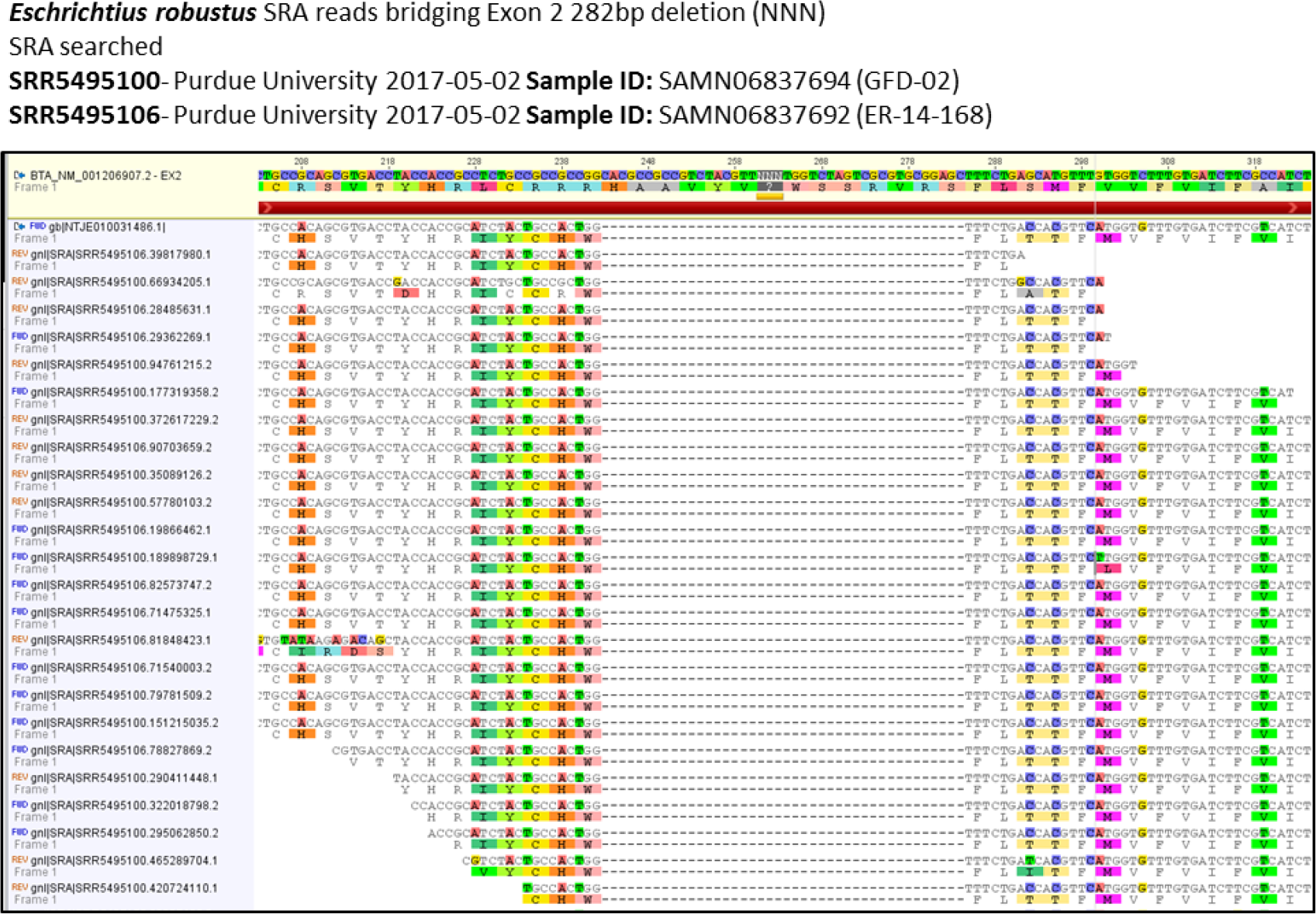
SRA validation for nucleotide deletion in exon 2 of *Mtnr1b* from *E. robustus*.

**Supplementary Figure 21:**
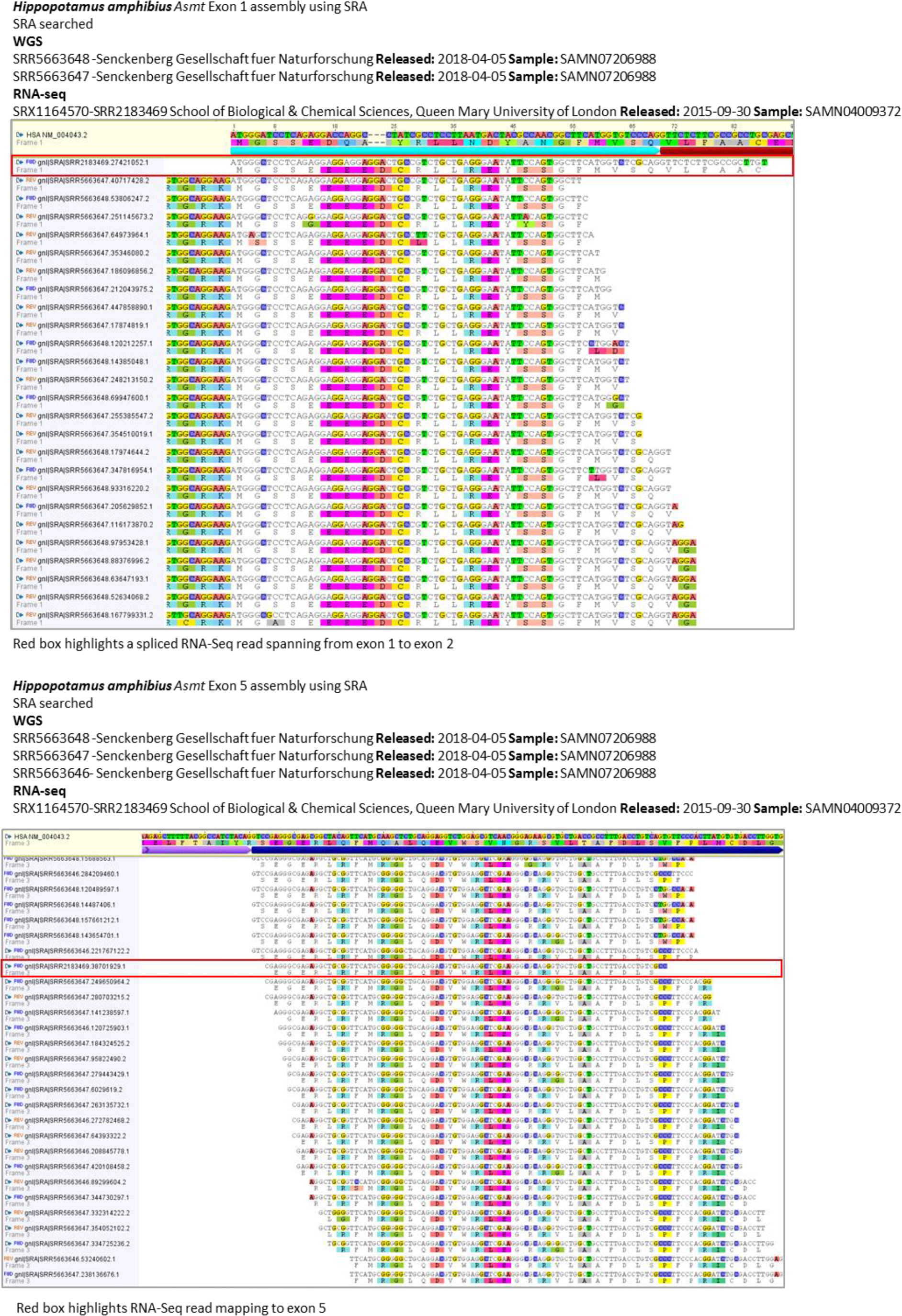
*H. amphibius Asmt* exon 1 and 5 assembly.

**Supplementary Figure 22:**
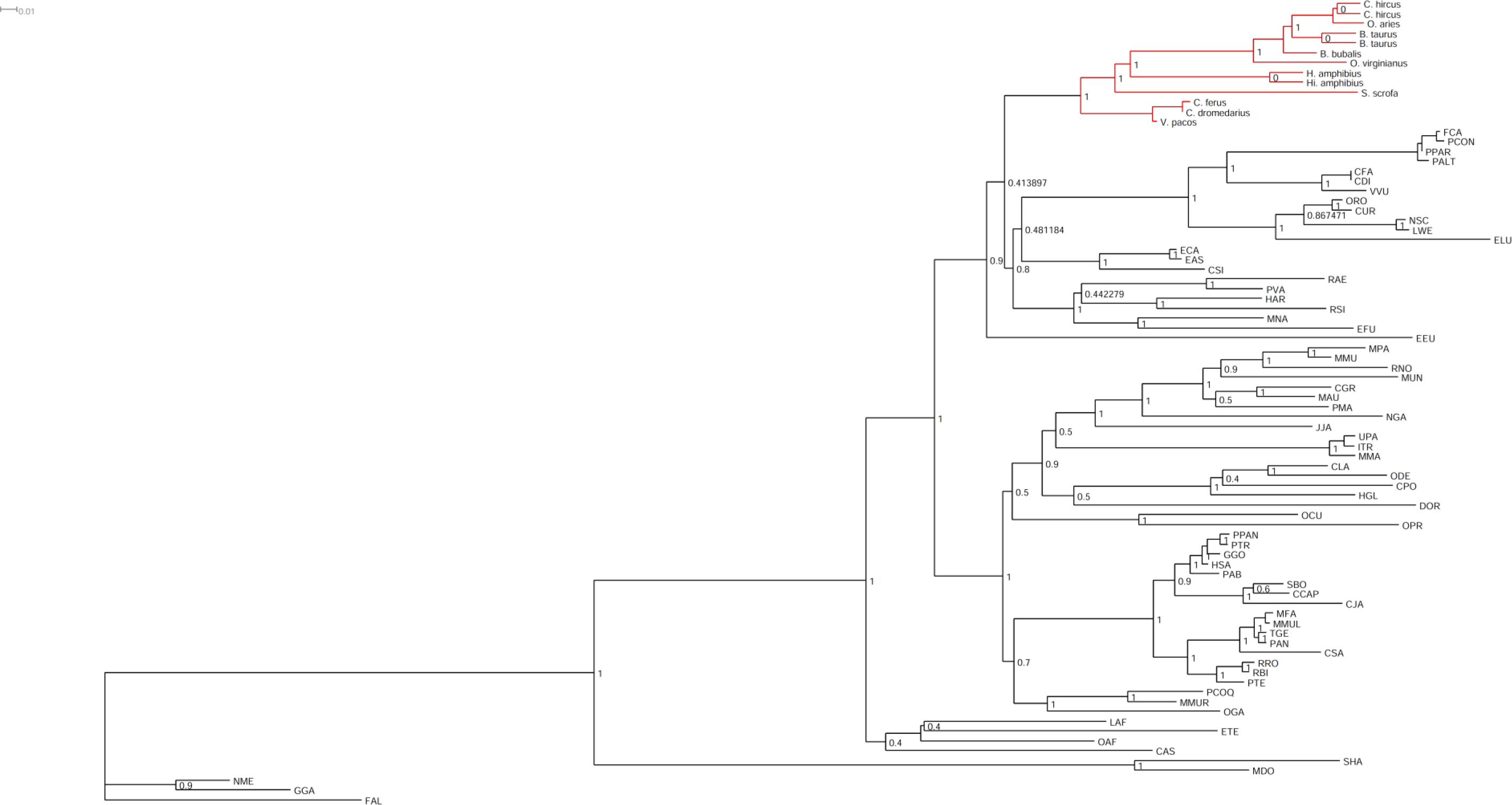
Maximum likelihood phylogenetic analysis of mammalian *Aanat* genes. Node values represent branch support using the aBayes algorithm. In red the clade containing the *H. amphibius*.

**Supplementary Figure 23:**
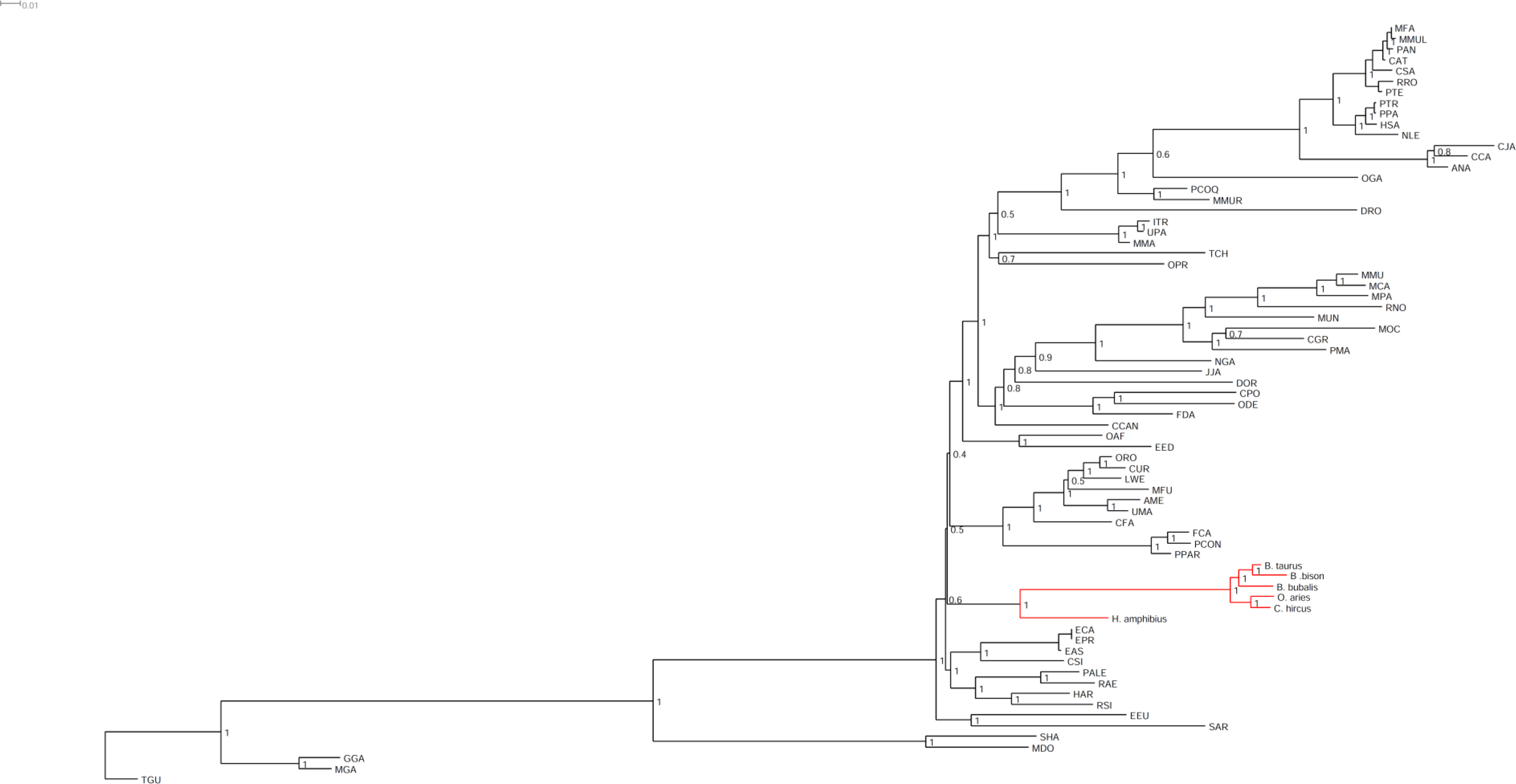
Maximum likelihood phylogenetic analysis of mammalian *Mtnr1a* genes. Node values represent branch support using the aBayes algorithm. In red the clade containing the *H. amphibius*.

**Supplementary Figure 24:**
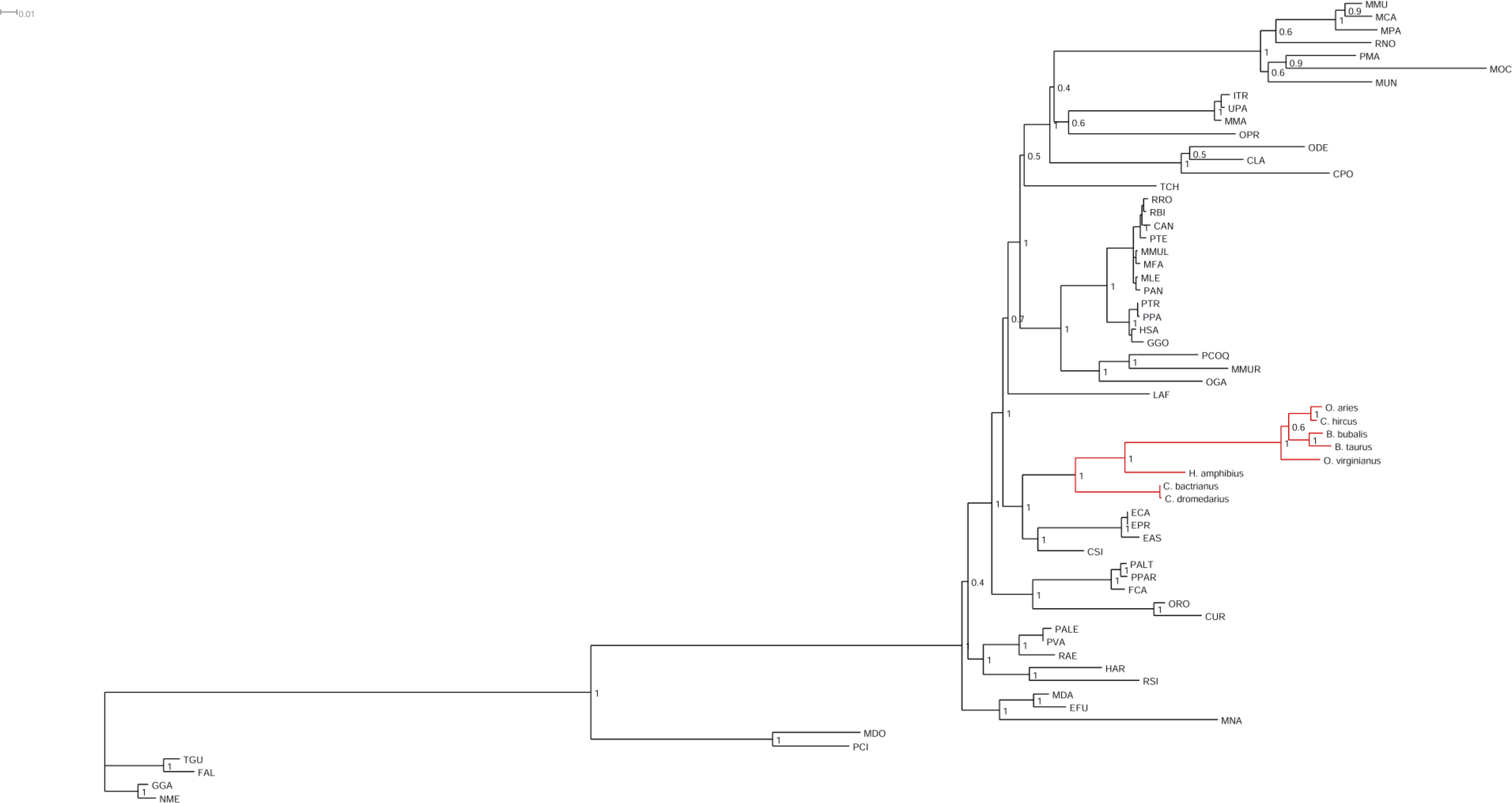
Maximum likelihood phylogenetic analysis of mammalian *Mtnr1b* genes. Node values represent branch support using the aBayes algorithm. In red the clade containing the *H. amphibius*.

**Supplementary Table 1:** Blast-n output of *Bos taurus* and *Homo sapiens* sequences against the *Hippopotamus amphibius* genome. These values were used to find the loci of *Hippopotamus amphibius* that contains the genes *Aanat*, *Mtnr1a* and *Mtnr1b*.

| Species | Accession numbers | Gene symbol | Scaffolds of *H. amphibius* genome | ID% | Exon number | Length | Gaps | qstart | qend | sstart | send | E-value | Bitscore |
| --- | --- | --- | --- | --- | --- | --- | --- | --- | --- | --- | --- | --- | --- |
| *Bos taurus* | NM_177509.2 | AANAT | NKPW01005924.1 | 87.117 | 1 | 163 | 0 | 1 | 163 | 56196 | 56034 | 1.71e-44 | 185 |
| *Bos taurus* | NM_177509.2 | AANAT | NKPW01005924.1 | 91.720 | 2 | 157 | 0 | 163 | 319 | 55714 | 55558 | 1.68e-54 | 219 |
| *Bos taurus* | NM_177509.2 | AANAT | NKPW01005924.1 | 86.452 | 3 | 310 | 2 | 316 | 624 | 55168 | 54860 | 1.24e-90 | 339 |
| *Bos taurus* | XM_024981048.1 | AANAT | NKPW01005924.1 | 87.730 | 1 | 163 | 0 | 1 | 163 | 44541 | 44379 | 3.66e-46 | 191 |
| *Bos taurus* | XM_024981048.1 | AANAT | NKPW01005924.1 | 90.604 | 2 | 149 | 0 | 163 | 311 | 44059 | 43911 | 2.19e-48 | 198 |
| *Bos taurus* | XM_024981048.1 | AANAT | NKPW01005924.1 | 83.495 | 3 | 309 | 0 | 316 | 624 | 43518 | 43210 | 1.26e-75 | 289 |
| *Bos taurus* | NM_001206907.2 | MTNR1B | NKPW01013551.1 | 94.667 | 1 | 225 | 0 | 1 | 225 | 40814 | 40590 | 1.06e-93 | 350 |
| *Bos taurus* | NM_001206907.2 | MTNR1B | NKPW01013551.1 | 88.575 | 2 | 814 | 0 | 221 | 1034 | 26808 | 25995 | 0.0 | 989 |
| *Bos taurus* | NM_005958.4 | MTNR1A | NKPW01072562.1 | 84.737 | 1 | 190 | 6 | 5 | 185 | 789 | 978 | 3.79e-43 | 182 |
| *Bos taurus* | NM_005958.4 | MTNR1A | NKPW01009466.1 | 85.682 | 2 | 873 | 3 | 183 | 1053 | 13832 | 14702 | 0.0 | 917 |
| *Homo sapiens* | NM_004043.2 | ASMT | NKPW01039339.1 | 81.250 | 2 | 176 | 2 | 68 | 241 | 2180 | 2005 | 6.87E-29 | 141 |
| *Homo sapiens* | NM_004043.2 | ASMT | NKPW01075055.1 | 78.333 | 5 | 120 | 0 | 441 | 560 | 1215 | 1334 | 5.46E-10 | 78.7 |
| *Homo sapiens* | NM_004043.2 | ASMT | NKPW01040565.1 | 77.778 | 7 | 144 | 1 | 653 | 795 | 6591 | 6734 | 9.08E-13 | 87.9 |
| *Homo sapiens* | NM_004043.2 | ASMT | NKPW01040565.1 | 86.667 | 8 | 135 | 3 | 786 | 918 | 8397 | 8530 | 1.48E-30 | 147 |
| *Homo sapiens* | NM_004043.2 | ASMT | NKPW01040565.1 | 74.528 | 9 | 212 | 2 | 908 | 1118 | 2399 | 2609 | 7.02E-14 | 91.6 |

**Supplementary Table 2:**
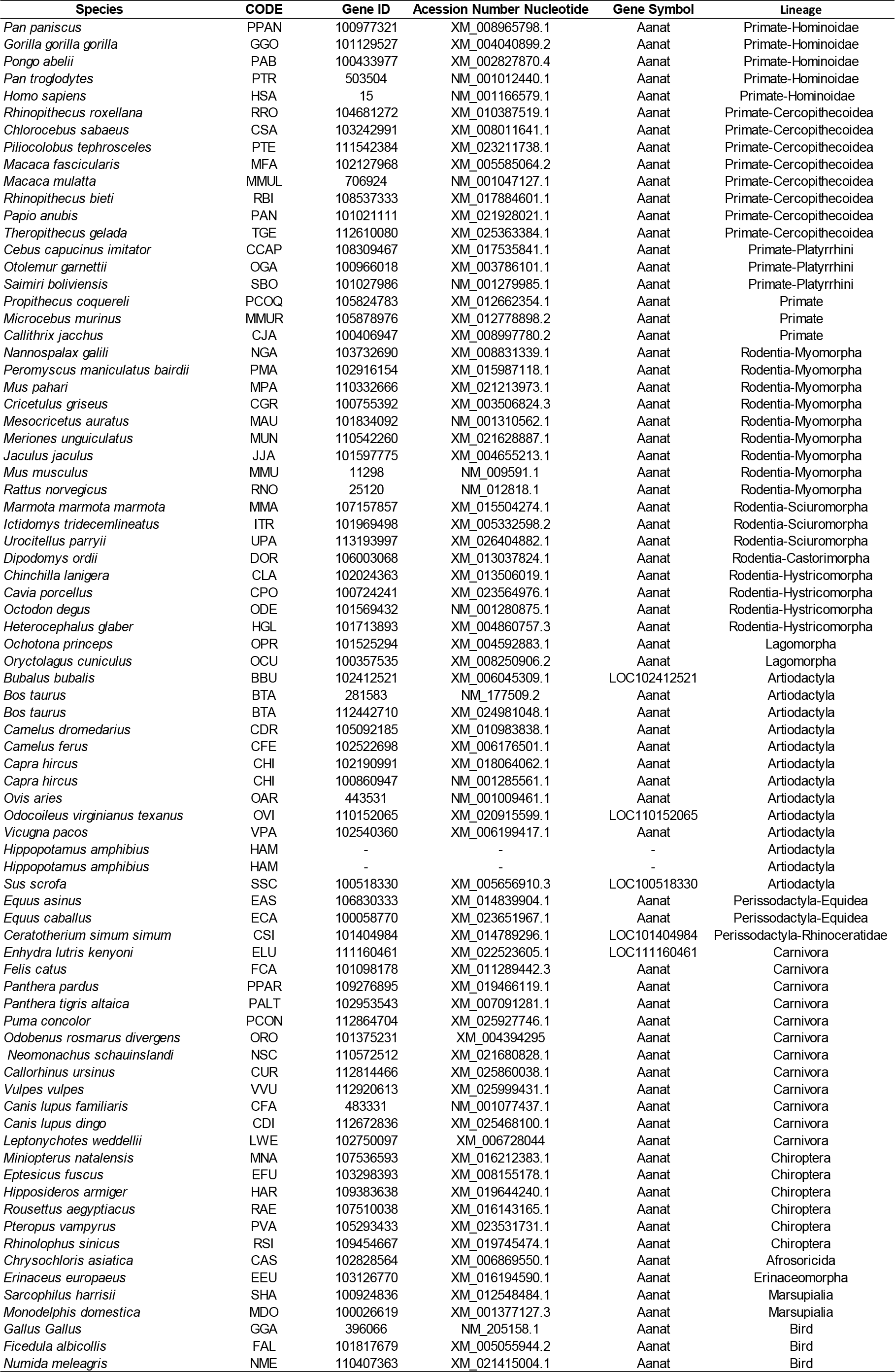
Accession numbers of the *Aanat* orthologues sequences used in phylogenetic analyses.

**Supplementary Table 3:**
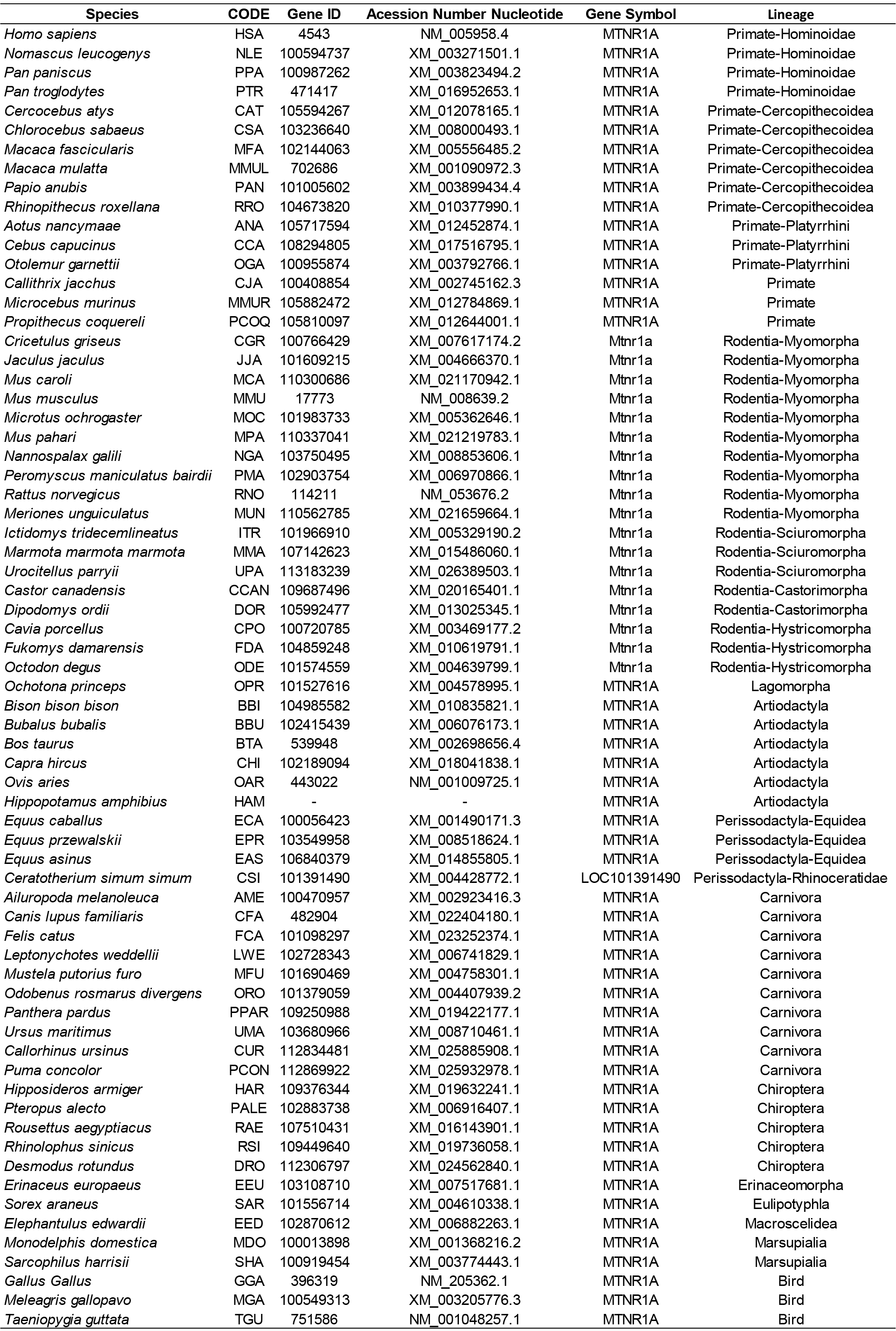
Accession numbers of the *Mtnr1a* orthologues sequences used in phylogenetic analyses.

**Supplementary Table 4:**
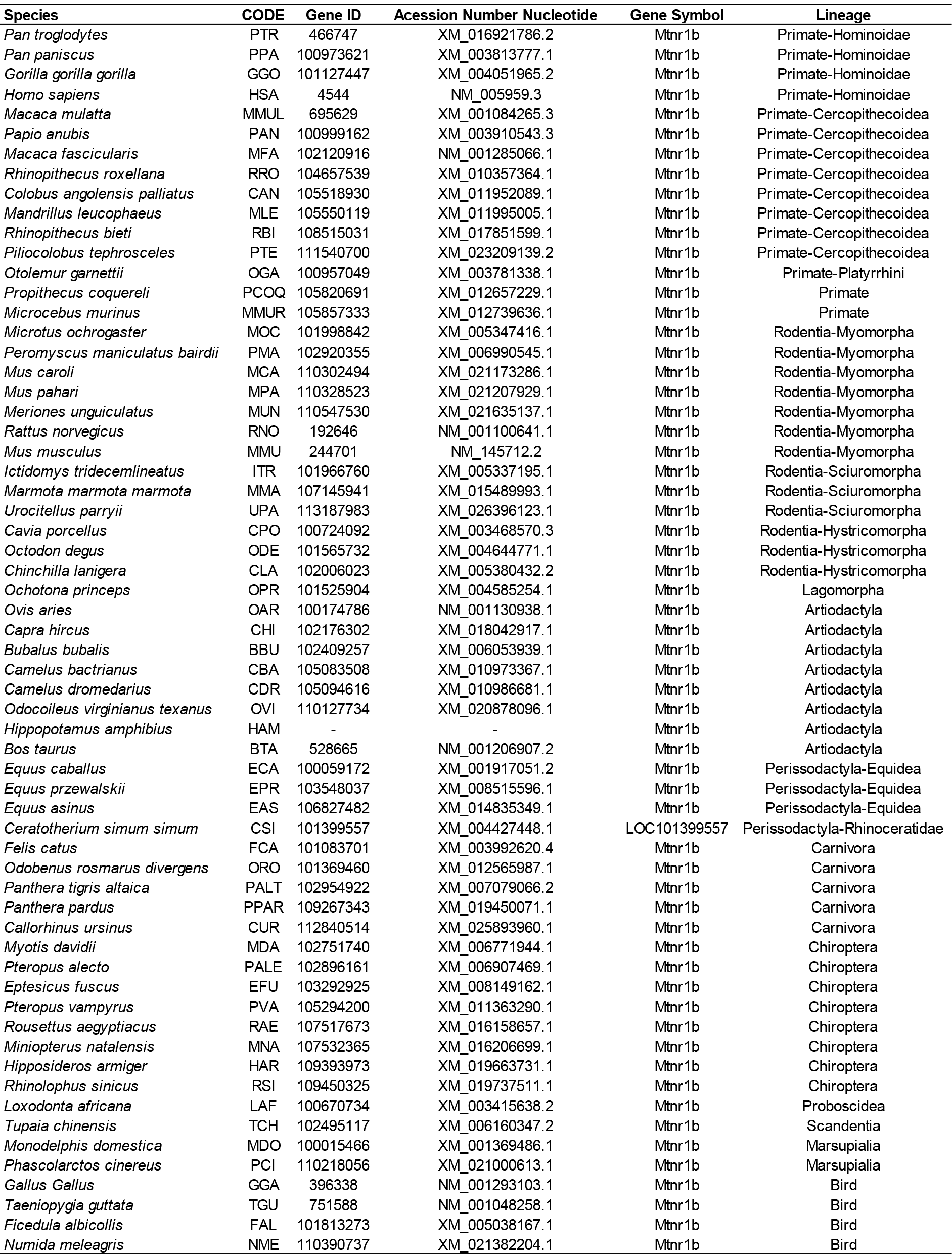
Accession numbers of the *Mtnr1b* orthologues sequences used in phylogenetic analyses.

## Gene annotations

In silico gene annotations of *Aanat*, *Mtnr1a* and Mtnr1b in *H. amphibius*

### *Aanat* genes of Hippopotamus amphibius annotated by Augustus Software

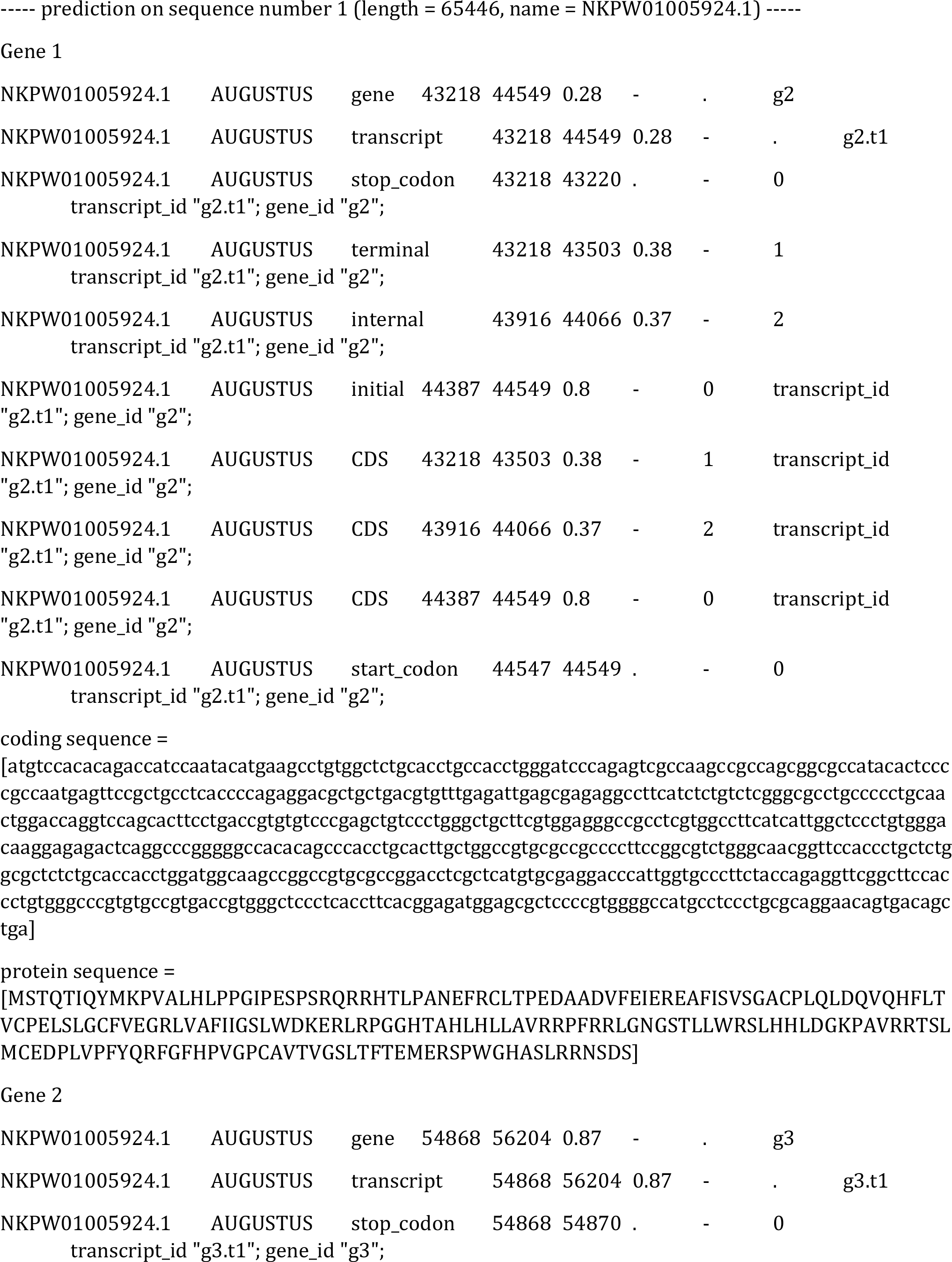

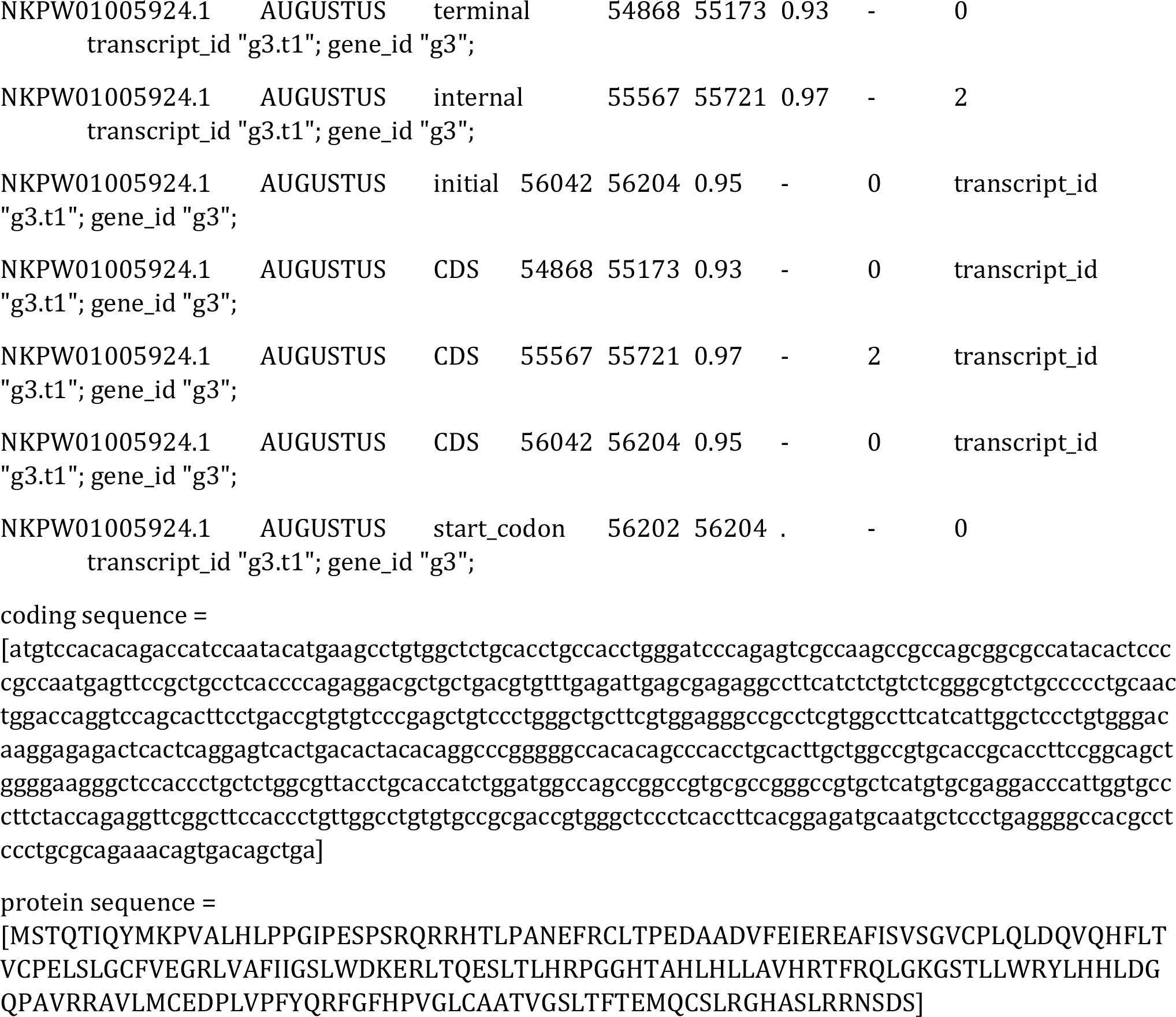

|  |  |  |  |  |  |  |  |  |
| --- | --- | --- | --- | --- | --- | --- | --- | --- |
| NKPW01005924.1 | AUGUSTUS | gene | 43218 | 44549 | 0.28 | - | . | g2 |
| NKPW01005924.1 | AUGUSTUS | transcript | 43218 | 44549 | 0.28 | - | . | g2.t1 |
| NKPW01005924.1 | AUGUSTUS | stop_codon | 43218 | 43220 | . | - | 0 |  |
| transcript_id "g2.t1"; gene_id "g2"; |  |  |  |  |  |  |  |  |
| NKPW01005924.1 | AUGUSTUS | terminal | 43218 | 43503 | 0.38 | - | 1 |  |
| transcript_id "g2.t1"; gene_id "g2"; |  |  |  |  |  |  |  |  |
| NKPW01005924.1 | AUGUSTUS | internal | 43916 | 44066 | 0.37 | - | 2 |  |
| transcript_id "g2.t1"; gene_id "g2"; |  |  |  |  |  |  |  |  |
| NKPW01005924.1 | AUGUSTUS | initial | 44387 | 44549 | 0.8 | - | 0 | transcript_id |
| "g2.t1"; gene_id "g2"; |  |  |  |  |  |  |  |  |
| NKPW01005924.1 | AUGUSTUS | CDS | 43218 | 43503 | 0.38 | - | 1 | transcript_id |
| "g2.t1"; gene_id "g2"; |  |  |  |  |  |  |  |  |
| NKPW01005924.1 | AUGUSTUS | CDS | 43916 | 44066 | 0.37 | - | 2 | transcript_id |
| "g2.t1"; gene_id "g2"; |  |  |  |  |  |  |  |  |
| NKPW01005924.1 | AUGUSTUS | CDS | 44387 | 44549 | 0.8 | - | 0 | transcript_id |
| "g2.t1"; gene_id "g2"; |  |  |  |  |  |  |  |  |
| NKPW01005924.1 | AUGUSTUS | start_codon | 44547 | 44549 | . | - | 0 |  |
| transcript_id "g2.t1"; gene_id "g2"; |  |  |  |  |  |  |  |  |
| NKPW01005924.1 | AUGUSTUS | gene | 54868 | 56204 | 0.87 | - | . | g3 |
| NKPW01005924.1 | AUGUSTUS | transcript | 54868 | 56204 | 0.87 | - | . | g3.t1 |
| NKPW01005924.1 | AUGUSTUS | stop_codon | 54868 | 54870 | . | - | 0 |  |
| transcript_id "g3.t1"; gene_id "g3"; |  |  |  |  |  |  |  |  |
| NKPW01005924.1 | AUGUSTUS | terminal | 54868 | 55173 | 0.93 | - | 0 |  |
| transcript_id "g3.t1"; gene_id "g3"; |  |  |  |  |  |  |  |  |
| NKPW01005924.1 | AUGUSTUS | internal | 55567 | 55721 | 0.97 | - | 2 |  |
| transcript_id "g3.t1"; gene_id "g3"; |  |  |  |  |  |  |  |  |
| NKPW01005924.1 | AUGUSTUS | initial | 56042 | 56204 | 0.95 | - | 0 | transcript_id |
| "g3.t1"; gene_id "g3"; |  |  |  |  |  |  |  |  |
| NKPW01005924.1 | AUGUSTUS | CDS | 54868 | 55173 | 0.93 | - | 0 | transcript_id |
| "g3.t1"; gene_id "g3"; |  |  |  |  |  |  |  |  |
| NKPW01005924.1 | AUGUSTUS | CDS | 55567 | 55721 | 0.97 | - | 2 | transcript_id |
| "g3.t1"; gene_id "g3"; |  |  |  |  |  |  |  |  |
| NKPW01005924.1 | AUGUSTUS | CDS | 56042 | 56204 | 0.95 | - | 0 | transcript_id |
| "g3.t1"; gene_id "g3"; |  |  |  |  |  |  |  |  |
| NKPW01005924.1 | AUGUSTUS | start_codon | 56202 | 56204 | . | - | 0 |  |
| transcript_id "g3.t1"; gene_id "g3"; |  |  |  |  |  |  |  |  |

coding sequence =
protein sequence =
``` [MSTQTIQYMKPVALHLPPGIPESPSRQRRHTLPANEFRCCLTPEDAADVFEIEREAFISVSGVCPLQLDQVQHFLT VCPELSLGCFFVEGRLVAFIIGSLWDERLTQESLTLHRPGGHTAHLHLLAVHRTFRQLGKGSTLLWRYLHHLDG QPAVRRRAVLNMCEDPLVPFYQRFQFHPVGLCAATVGLSLTFTEMQCSLRGHASLRNDS] ```

### *Mtnr1a* gene of Hippopotamus amphibius annotated by Augustus Software

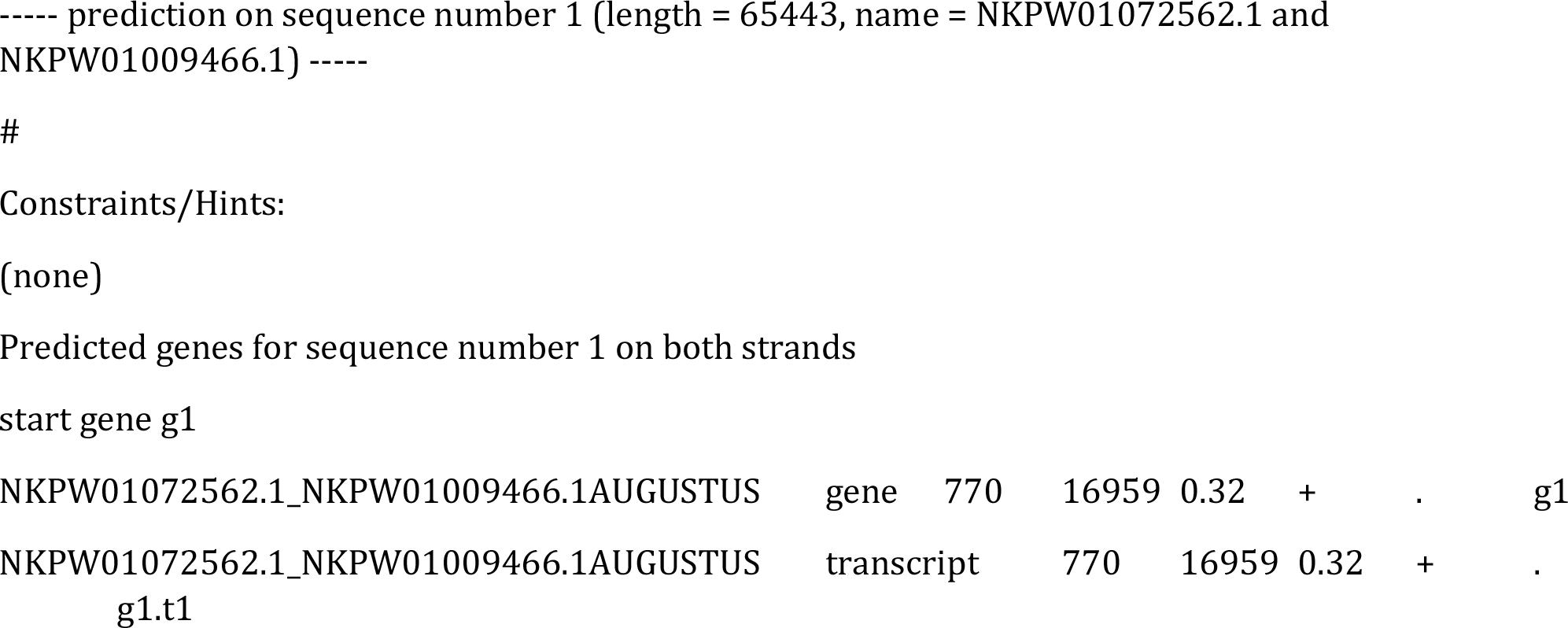

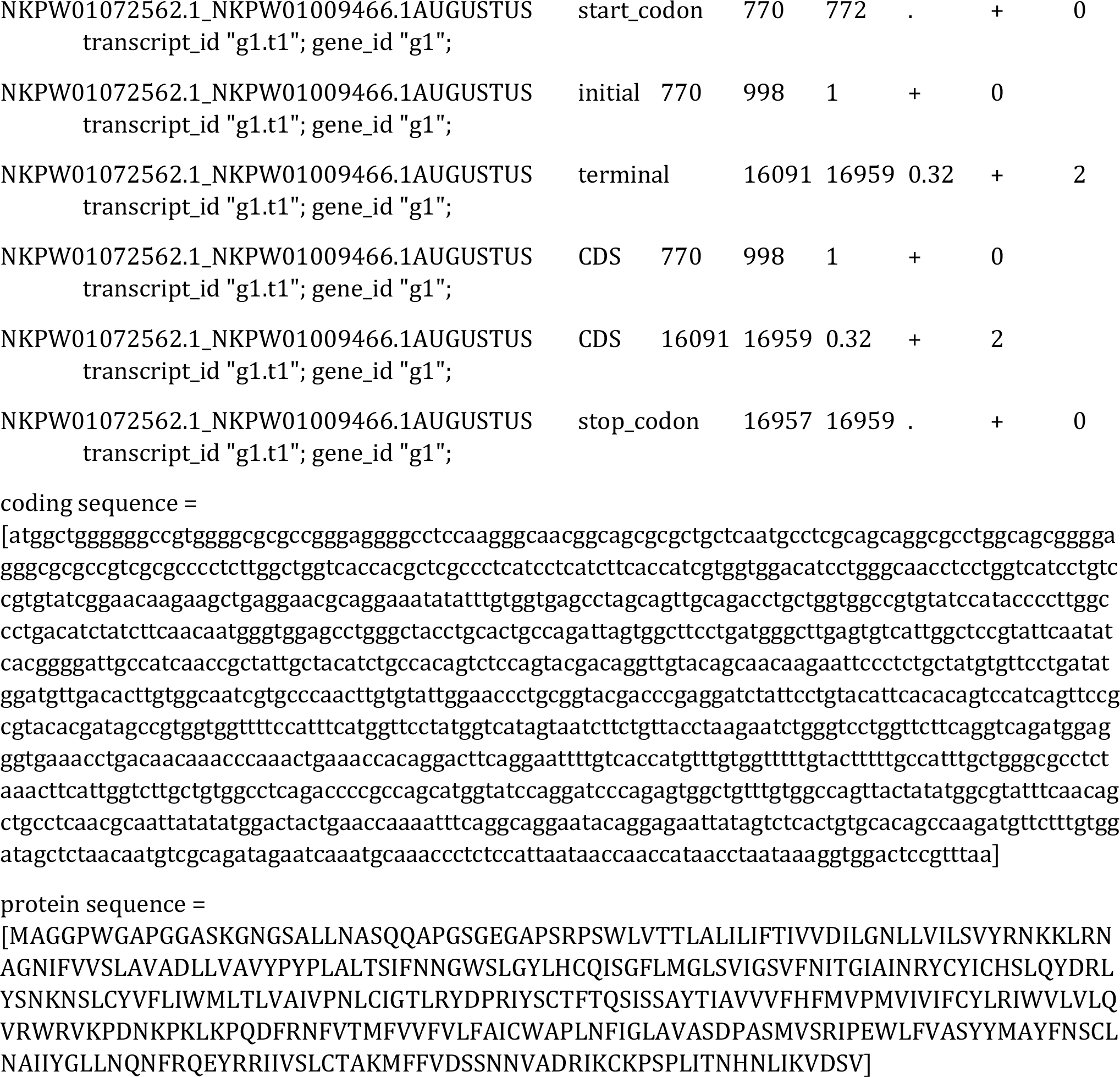

|  |  |  |  |  |  |  |  |
| --- | --- | --- | --- | --- | --- | --- | --- |
| NKPW01072562.1_NKPW01009466.1AUGUSTUS | gene | 770 | 16959 | 0.32 | + | . | g1 |
| NKPW01072562.1_NKPW01009466.1AUGUSTUS | transcript | 770 | 16959 | 0.32 | + | . |  |
| g1.t1 |  |  |  |  |  |  |  |

|  |  |  |  |  |  |  |
| --- | --- | --- | --- | --- | --- | --- |
| NKPW01072562.1_NKPW01009466.1AUGUSTUS | start_codon | 770 | 772 | . | + | 0 |
| transcript_id "g1.t1"; gene_id "g1"; |  |  |  |  |  |  |
| NKPW01072562.1_NKPW01009466.1AUGUSTUS | initial | 770 | 998 | 1 | + | 0 |
| transcript_id "g1.t1"; gene_id "g1"; |  |  |  |  |  |  |
| NKPW01072562.1_NKPW01009466.1AUGUSTUS | terminal | 16091 | 16959 | 0.32 | + | 2 |
| transcript_id "g1.t1"; gene_id "g1"; |  |  |  |  |  |  |
| NKPW01072562.1_NKPW01009466.1AUGUSTUS | CDS | 770 | 998 | 1 | + | 0 |
| transcript_id "g1.t1"; gene_id "g1"; |  |  |  |  |  |  |
| NKPW01072562.1_NKPW01009466.1AUGUSTUS | CDS | 16091 | 16959 | 0.32 | + | 2 |
| transcript_id "g1.t1"; gene_id "g1"; |  |  |  |  |  |  |
| NKPW01072562.1_NKPW01009466.1AUGUSTUS | stop_codon | 16957 | 16959 | . | + | 0 |
| transcript_id "g1.t1"; gene_id "g1"; |  |  |  |  |  |  |

coding sequence =
protein sequence =
``` [MAGGPWGAPGGASKGNGSALLNASQQAPGSGEGAPSRPSWLVTTLALILIFTIVVDILGNLLVILSVYRNKKLRN AGNIFVVS LAVADLLVAVYPYPLALTSIFNNGWSLGYLHCQISGFLMGLSVIGSVFNITGIAINRYCYICHSLQYDRL YSNKNSLCYVFLIWMLTLVAIVPNLCIGTLRYDPRIYSCTFTQSISSAYTIAVVVFHFMVPMVIVIFCYLRIWVLVLQ VRWRVKPDNPKPKLPQDFRNFVTMFVVFVLFALCWAPLNFIFGLAVASDPASMVSRIPEWLFVASYMAYFNSCL NAIYGLLNQNFRQEYRRIIVSLCTAKMFFVDSSNNVADRIKCKPSPLITNHNLIKVDSV] ```

### *Mtnr1b* gene of Hippopotamus amphibius annotated by Augustus Software

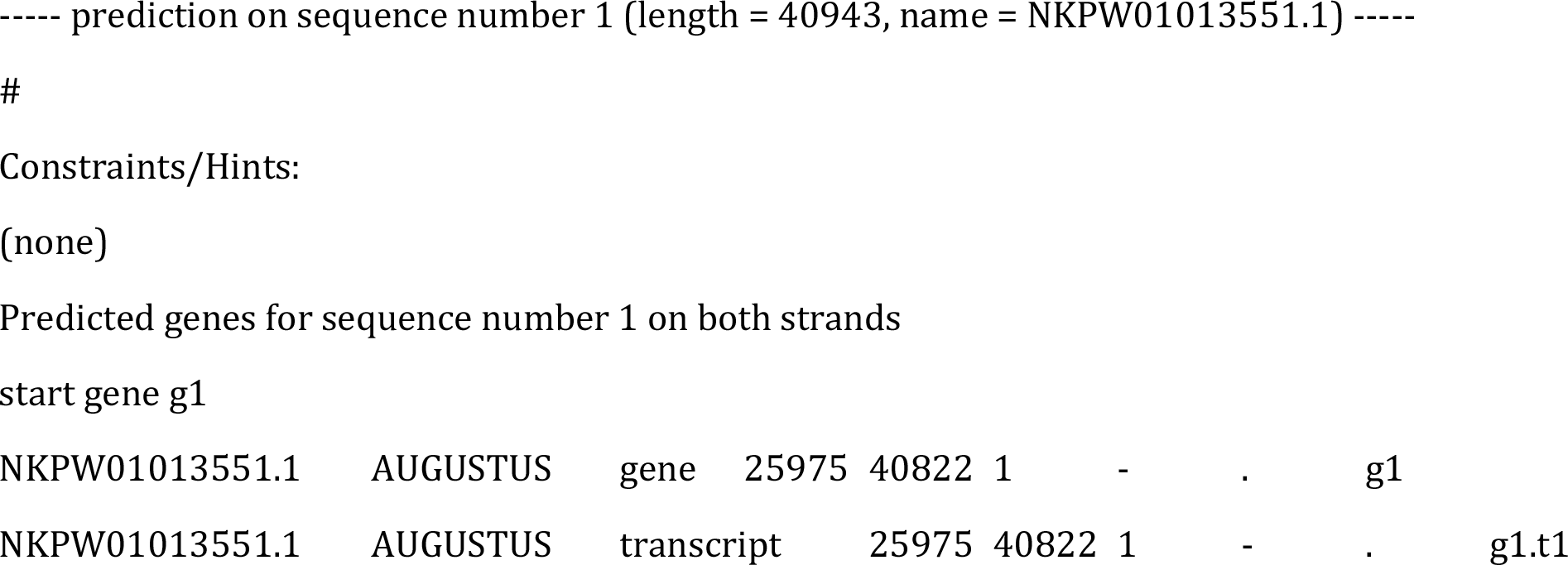

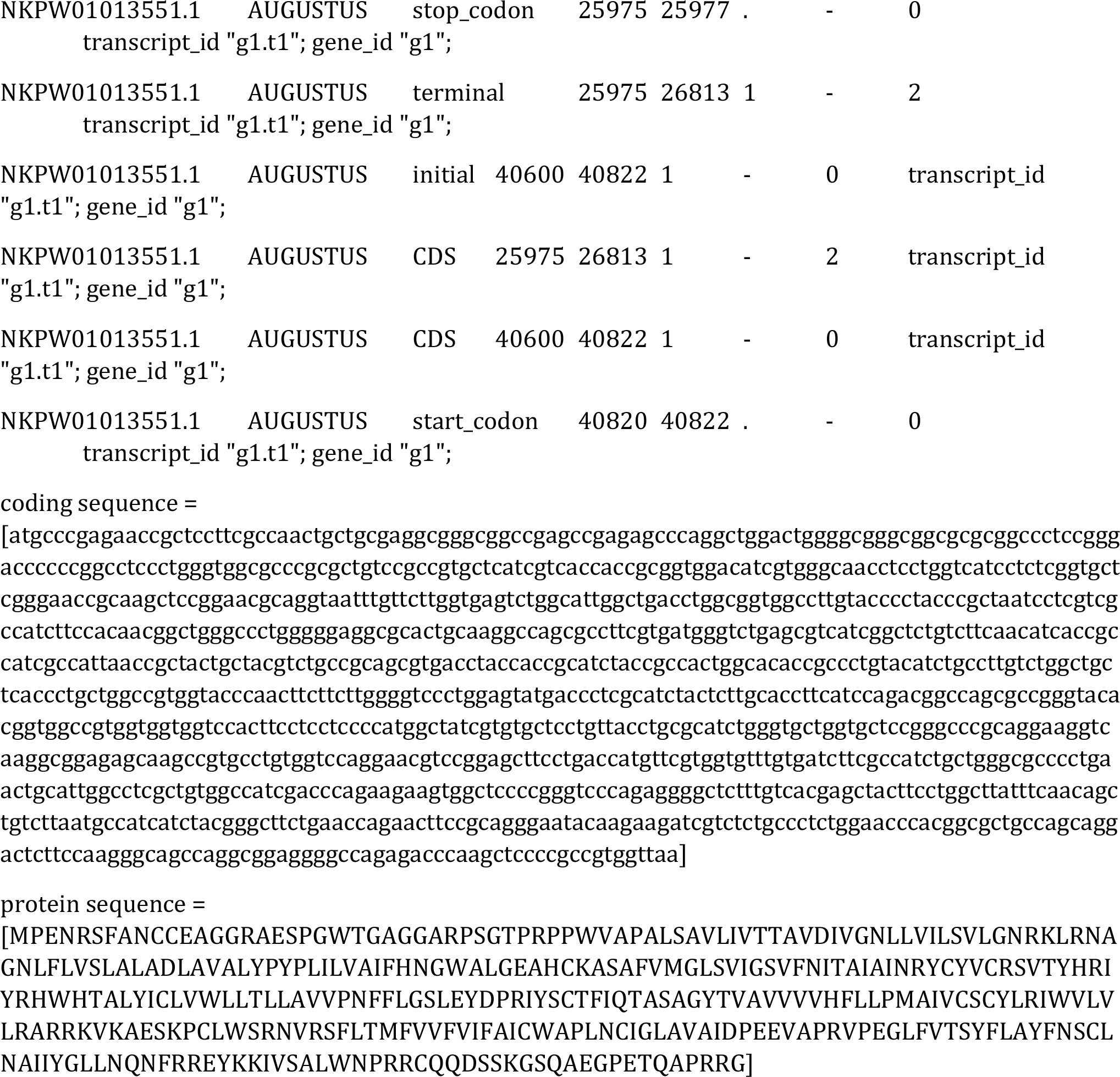

|  |  |  |  |  |  |  |  |  |
| --- | --- | --- | --- | --- | --- | --- | --- | --- |
| NKPW01013551.1 | AUGUSTUS | gene | 25975 | 40822 | 1 | - | . | g1 |
| NKPW01013551.1 | AUGUSTUS | transcript | 25975 | 40822 | 1 | - | . | g1.t1 |
| NKPW01013551.1 | AUGUSTUS | stop_codon | 25975 | 25977 | . | - | 0 |  |
| transcript_id "g1.t1"; gene_id "g1"; |  |  |  |  |  |  |  |  |
| NKPW01013551.1 | AUGUSTUS | terminal | 25975 | 26813 | 1 | - | 2 |  |
| transcript_id "g1.t1"; gene_id "g1"; |  |  |  |  |  |  |  |  |
| NKPW01013551.1 | AUGUSTUS | initial | 40600 | 40822 | 1 | - | 0 | transcript_id |
| "g1.t1"; gene_id "g1"; |  |  |  |  |  |  |  |  |
| NKPW01013551.1 | AUGUSTUS | CDS | 25975 | 26813 | 1 | - | 2 | transcript_id |
| "g1.t1"; gene_id "g1"; |  |  |  |  |  |  |  |  |
| NKPW01013551.1 | AUGUSTUS | CDS | 40600 | 40822 | 1 | - | 0 | transcript_id |
| "g1.t1"; gene_id "g1"; |  |  |  |  |  |  |  |  |
| NKPW01013551.1 | AUGUSTUS | start_codon | 40820 | 40822 | . | - | 0 |  |
| transcript_id "g1.t1"; gene_id "g1"; |  |  |  |  |  |  |  |  |

coding sequence =
protein sequence =
``` [MPENRSFANCCEAGGRAESPGWTGAGGARPSGTPRPPWVAPALSAVLIVTTAVDIVGNLLVILSVLGNRKLRNA GNLFLVSLALADLAVALYPYPLILVAIFHNGWALGEAHCKASAFVFMGLSVIGSVFNITAIINRYCYVCRSVTYHRI YRHWHTALYICLVWLLTLLAVVPNFFLGSLEYDPRIYSCTFIQTASAGYTVAVVVVHFLPMAIVCSCYLRIWVLV LRARRKVKAESKPCLWSRNVRSLTMFVVFVIFAICWAPLNCIGLAVDAIDPEEVAPRVPEGLFVTSYFLAYFNSCL NAIYGLLNQNFREYKKIVSALWNP RRCCQDSSKGSQAEGPETQAPRRG] ```

